# Beyond additivity: zero-shot methods cannot predict impact of epistasis on protein properties and function

**DOI:** 10.64898/2026.02.17.706292

**Authors:** Anastasia Kolchina, Igors Dubanevics, Fyodor A. Kondrashov, Olga V. Kalinina

## Abstract

Accurate prediction of properties and function of mutated proteins is crucial for both research and industrial applications. Experimental assessment of mutations relies on biochemical techniques, which, while accurate, are costly and labour-intensive. As an alternative, computational methods have emerged as a scalable and cost-effective solution. A key challenge for predicting functional consequences of mutations is epistasis, a phenomenon where the effect of one mutation is influenced by others. We evaluated the ability of 95 zero-shot models to predict the impact of epistasis on proteins using datasets from ProteinGym. Our results demonstrate that while the current models perform well for single mutations and non-epistatic combinations of mutations, they fail to predict the effect of strongly epistatic combinations of mutations. This exposes deficiencies of the state-of-the-art models and the need for focusing on capturing complex mutational interactions, which is essential for advancing both evolutionary studies and protein design.

## Main

The impact of a mutation on protein properties and function depends on the presence of other mutations in that protein: an effect known as epistasis. Intragenic epistasis is omnipresent^1^, whereas the extent of intergenic epistasis is less well understood but thought to be comparably rare^2^. Physical interactions between amino acid residues in protein structure is an intuitively obvious functional basis of epistatic interactions^3,4^. However, large scale experiments show that physical interactions explain only a fraction of all intergenic epistatic interactions, and hence predicting epistatic effects cannot be reduced to predicting the protein structure alone^5^. This makes *in silico* prediction of the genotype-to-phenotype relationships defined by epistasis a yet-to-be-solved problem.

Most mutations are either neutral or have a slightly to severely detrimental effect for protein function, thus, in an unchanging environment wildtype gene sequences occupy fitness peaks in the vast fitness landscape that spans the entire sequence space^6^ (Figure 1a). Yet the fitness landscapes of many proteins are rugged, meaning that they have multiple isolated fitness peaks^5,7–11^. Such rugged topology must be caused by epistatic interactions of different mutations^12^ and would be impossible if protein function that defines fitness could be determined by a simple sum or a linear combination of effects of individual variants (Figure 1a). Epistatic interactions are essential for many protein properties and functions, including binding affinity^7^, stability^13^, and enzyme activity^14^.

**Figure 1.**
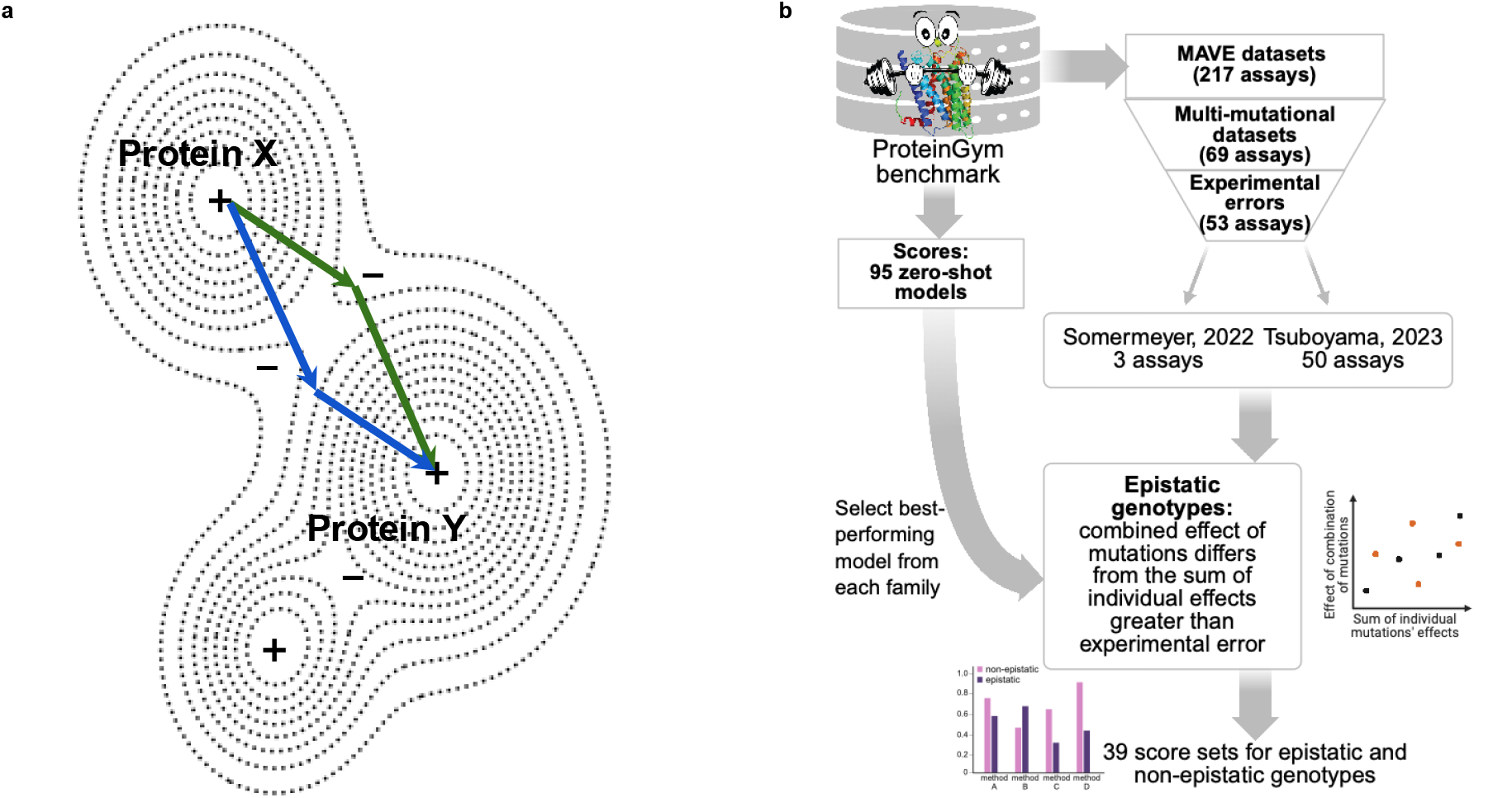
**a**. A sketch of a fitness landscape (after Wright 1932^24^). Wildtype sequences reside at fitness peaks, denoted with a ‘+’ sign, which are separated by valleys of low fitness, denoted with a ‘-’ sign; dotted lines separate iso-fitness areas. Possible mutational paths between Protein X and Protein Y are shown with green and blue chains of arrows; all such paths require traversing a valley of low fitness. Hence, epistasis is required to travel from one fitness peak to another. **b**. Analysis pipeline: from the MAVE datasets in ProteinGym^23^, we selected those where data on experimental error were available and identified epistatic genotypes within them. We assessed performance of zero-shot methods specifically for these genotypes, and from each family of methods (for example, similar models only differing by the number of parameters) selected the best-performing one.

Even for small proteins, if one considers the interaction of a large number of mutations, the number of possible sequence variants is astronomically large and impossible to ascertain experimentally. Thus, there is a pressing need to develop computational tools that can address the problem of predicting the functional impact of multiple mutations that take into account epistatic genetic interaction effects. Variant effect prediction (VEP) methods can accurately predict the effect of a single mutation in an otherwise functional protein^15,16^. Recently, machine-learning and deep-learning methods are dominating the field, but they hardly perform better than simple statistical tools that rely on key features such as evolutionary conservation^16^. Moreover, most VEP methods have not been designed to predict the impact of several concurrent mutations^15^.

Machine- and deep-learning methods have performed exceptionally well in various complex tasks, in which the solution space is complex and driven by parameter interactions^17–19^. Among existing VEP methods are those that are trained on existing experimental data on protein activity (supervised methods) and those that forgo training on empirical data altogether (unsupervised or zero-shot methods). Supervised methods are inapplicable for studying epistasis. First, most of them are not designed to be generalizable to unseen proteins, as they are trained to predict the effect of mutations in a protein based on effects of other mutations in that same protein, and thus may exploit protein-specific labels, therefore not reflecting understanding of the protein’s mutational landscape. Second, the supervised methods are trained on existing experimental data, which typically comprise only information about the effect of single point mutations; hence, these supervised models can only predict the effect of one mutation at a time. Thus, by design of reliance on the limited empirical datasets, the supervised methods cannot be used to model larger sequence spaces.

Given the lack of empirical data available for model training, only zero-shot VEP approaches can be used to model changes of properties and function of sequences with multiple mutations. Such approaches are rising in popularity^15,16^. The zero-shot methods assess some notion of biological plausibility of a sequence from having seen many naturally occurring biological sequences. One example of such methods is the ESM family of protein language models^20^. These models were trained as masked language models, in that they learned to predict masked amino acids in a protein sequence using information about all protein sequences in the UniProt database^21^. In a naive setup such a model would simply learn sequence homology, so the authors of the ESM models took care that the training and test data would not contain too similar proteins.

To assess the utility of various models for variant effect prediction, large and diverse experimental data is provided by MAVE (multiplexed assays of variant effects) experiments. In these experiments, for a given protein the effect of (almost) every possible mutation at (almost) every position is measured^22^. Performance of VEP methods in these data is reported in ProteinGym^23^, which is the largest independent benchmark to date and comprises data on experimental effects of an almost exhaustive mutational scan of 217 different proteins or their regions derived from MAVE experiments, as well as data on clinical significance of over 66 thousand variants in the human genome.

In this study, we assess the extent of epistasis in the MAVE data and how well VEP tools predict the effect of combinations of epistatically interacting variants. We evaluated all 95 zero-shot models using the data from 53 MAVE datasets with measured effect values for multi-mutational variants from the ProteinGym^23^ benchmark (Figure 1b). We created simple baselines that only exploit information on the effect of single mutations and by design cannot capture epistasis. When considering combinations of the mutations with epistatic effects, we demonstrate that none of the VEP methods can predict these effects better than the baseline models. These results reveal a glaring lack of performant models for predicting non-linear epistatic effects. This calls for increasing efforts to generate more experimental data capturing the effects of variant combinations and to develop better computational tools to predict them.

## Results

### Overview of the dataset

ProteinGym reports an assessment of 95 zero-shot VEP methods for their ability to predict effects of variants in protein sequences. These methods by design are not trained on MAVE experimental data, but rather on unlabelled data (e.g. from the whole UniProt^21^), and thus predict some general notion of a variant’s fitness as it is represented by all available protein sequences.

Each dataset in ProteinGym comprises a set of variants – which we call *genotypes* – of a single protein – which we call *wildtype* – whose function and the effect of mutations on it have been experimentally assessed. Of the 217 MAVE datasets in ProteinGym, 69 contain experimental data for genotypes with more than one mutation, with some datasets containing as many as 44 mutations in a single genotype. As we wanted to take into account experimental uncertainty of measurements, we proceeded with the Somermeyer^11^ and Tsuboyama^25^ datasets, as information on experimental errors was available for them, which resulted in 53 datasets, three from the Somermeyer and 50 from the Tsuboyama studies, respectively (Figure 1b).

The Somermeyer datasets comprise three datasets for measured fluorescence values of mutants of three GFP proteins from *Aequorea macrodactyla* (amacGFP), *Clytia gregaria* (cgreGFP) and *Pontellina plumata* (ppluGFP), comprising 33,510, 24,515 and 31,401 genotypes, respectively. The distributions of the number of mutations per dataset are quite different in the three Somermeyer datasets (Figure 2a), with the dataset for *C. gregaria* and *P. plumata* containing many genotypes with two mutations from the wildtype, whereas the dataset for *A. macrodactyla* contains a largest fraction of three-mutation genotypes. This latter dataset contains also the largest number of genotypes with more than 10 mutations (43 vs. 23 and 3 for *C. gregaria* and *P. plumata*, respectively). The distribution of the fluorescence values is also not identical in the three datasets (Figure 2b).

**Figure 2.**
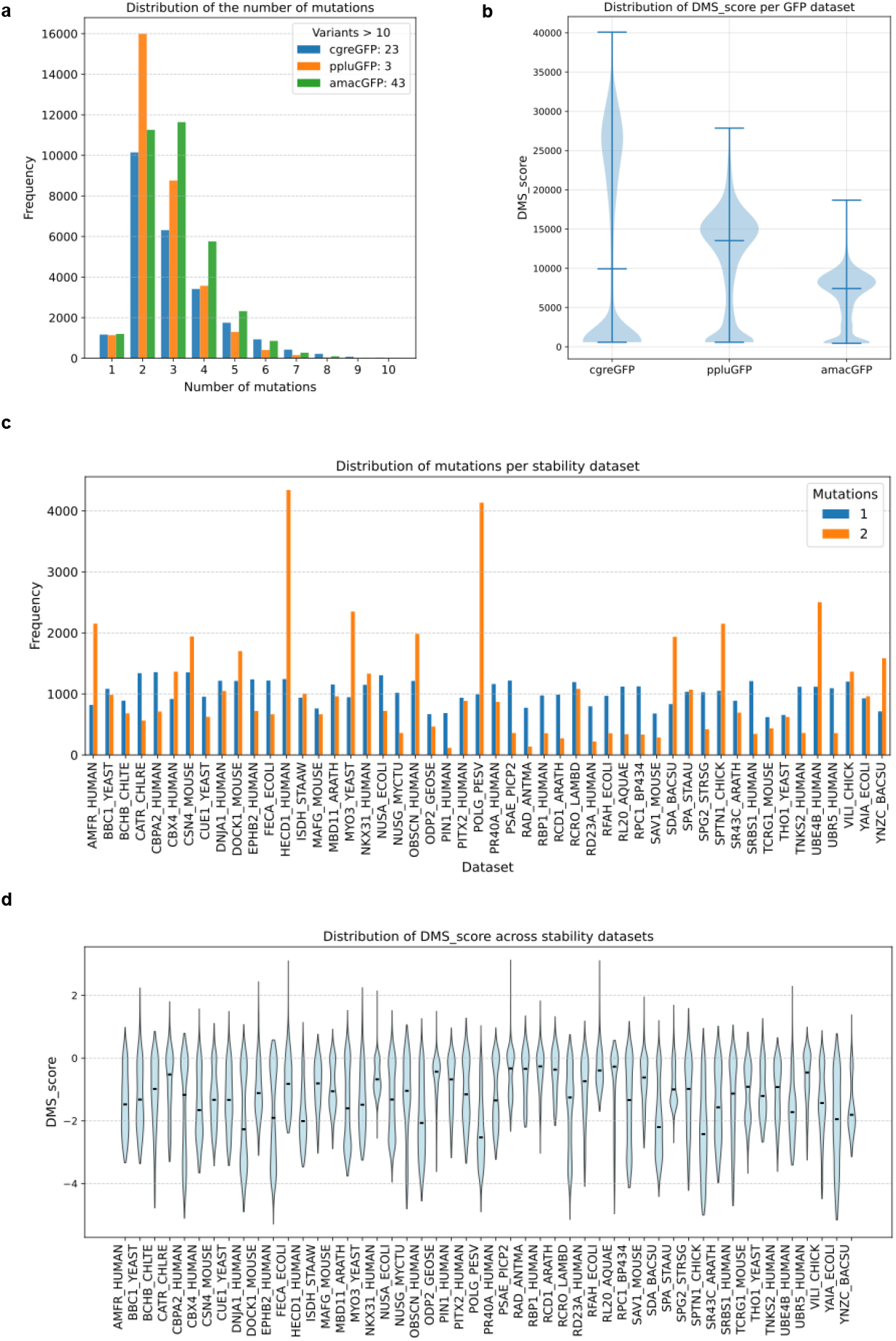
Mutation number and target value distributions. **a**. Distribution of mutation counts in the three GFP datasets (cgreGFP, ppluGFP, amacGFP; left to right). The x-axis shows the number of mutations per genotype, and the y-axis shows their frequency in the dataset. **b**. Distribution of mutation counts in the stability (Tsuboyama) datasets, each of which contains only single- and double-mutant genotypes. **c**. Distribution of MAVE scores (fluorescence values) for the GFP datasets. The histogram shows the frequencies of MAVE scores (x-axis) for each protein. **d**. Distribution of MAVE scores in the stability datasets. For each dataset on the x-axis, a box plot summarizes the corresponding thermostability values.

The Tsuboyama datasets contain fundamentally different information: measurements of thermostability for single and double mutants of 50 proteins of various functions, totalling 103,055 genotypes (between 802 and 5,586 genotypes per protein, Figure 2c). Despite the difference in length, function and the organism of origin, the change of thermostability is comparable across all datasets (Figure 2d).

### Magnitude of epistasis

For the GFP datasets, we define the magnitude of epistasis following the original study^11^:

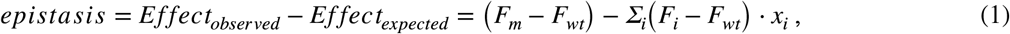

where *F*_*i*_, *F*_*m*_, *F*_*wt*_ are measured levels of effect of a genotype with a single mutation *i*, of genotype with several mutations, or of the wildtype sequence, respectively; *x*_*i*_ = 1 when mutation *i* is present in the genotype and *x*_*i*_ = 0 otherwise^11^. The higher the difference between the observed effect (experimental value for a multi-mutation genotype) and the expected effect (sum of experimental values for the corresponding single-mutation genotypes), the stronger the epistasis in the multi-mutation genotype (Figure 3a).

**Figure 3.**
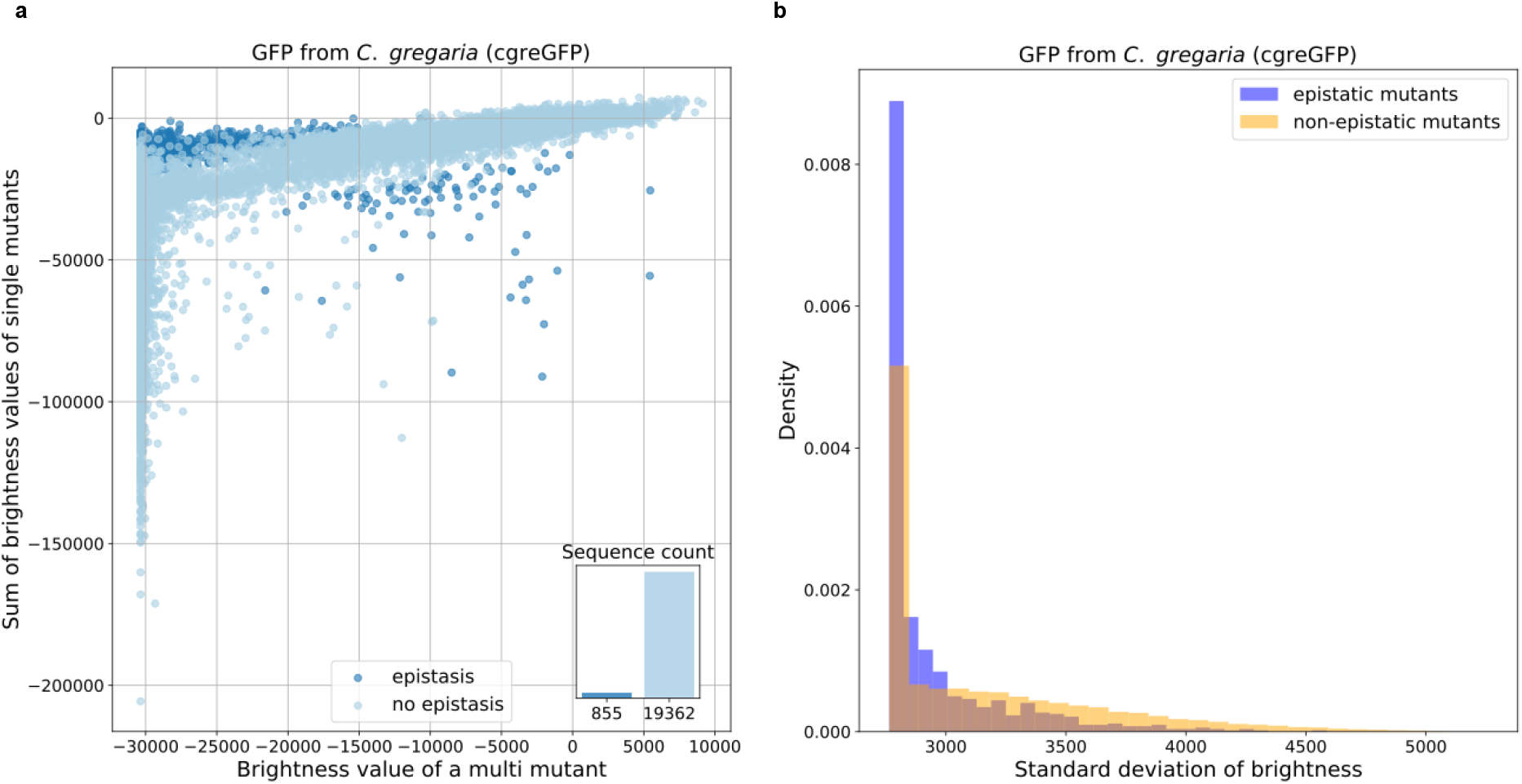
**a**. Epistatic and non-epistatic genotypes in the dataset for the GFP from *C. gregaria*^11^: for each multi-mutant genotype, the x-axis shows its observed brightness, and the y-axis shows the sum of the corresponding single-mutant brightness values. Both values are shown as raw brightness values (not log-transformed) and relative to the wildtype. Dark blue dots correspond to the epistatic genotypes, i.e. where the functional effect of a combination of mutations significantly (larger than the experimental error) differs from the linear combination of individual effects. **b**. The distribution of experimental errors in epistatic (dark blue) and non-epistatic (orange) genotypes for *C. gregaria*. Observed epistasis cannot be attributed to larger experimental errors and hence is a genuine biological effect. For other GFP datasets, see Supplementary Material.

For the stability datasets, instead of the sum of the effects of individual mutants, we used reconstructed *ΔG* values under the assumption of independence of contributions by individual mutations based on the thermodynamic model suggested by the authors of the original study (see Methods). We then selected epistatic sequences using a two-tailed Z-test to detect significance of epistasis:

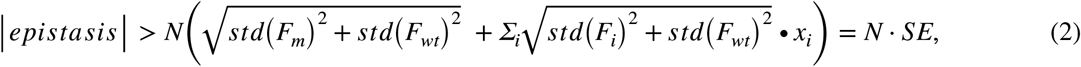

Where *std* is the standard deviation of the effect measurements equal to the reported experimental errors, which are either directly available for each dataset, or we extract them from the available data (see Methods). *SE* is the standard error of the estimated epistasis (Figure 3a).

*N* is a significance threshold expressed in std units added to detect strong epistatic effects; different values of *N* were explored in this study (Supplementary Figure 2). In the following, we chose *N* = 1 for the GFP and *N*= 3 for the stability datasets. This choice is motivated by the fact that the GFP datasets were specifically generated with the aim to study epistasis^11^ and contain a significant proportion of sequences with more than two mutations. They are also larger than the stability datasets in terms of the number of genotypes. The stability datasets contain at most two mutations, and each dataset is more than five-fold smaller than the GFP datasets. The relative magnitude of the standard deviation of the effect measurements is also larger in the stability datasets (Supplementary Figure 7). Additionally, the nature of the biological effect is different (fluorescence and thermostability) in the two cases, hence, selecting a common criterion for epistasis is probably impossible.

For the three GFP datasets, we identified 232, 859 and 959 epistatic genotypes for amacGFP, cgreGFP and ppluGFP respectively, and for the stability datasets, the number of epistatic genotypes lies between 4 and 645 (except one dataset with only one epistatic genotype, which we did not use further, Supplementary Table 1). It must be noted that the detected epistasis cannot be explained by higher experimental errors in these genotypes (Figure 3b): indeed, the distribution of experimental errors for the genotypes that we define as epistatic is even narrower than for non-epistatic ones. Hence, detected epistasis is a biologically relevant phenomenon and not an experimental artifact.

### Performance of zero-shot methods

We compared the agreement between fitness values predicted by 95 zero-shot methods and the experimental measurements for all tested genotypes and for epistatic genotypes separately. From each family of related methods (e.g. differing by only the number of parameters), we selected the best performing model and hence report 39 Spearman’s correlation values (Figures 4 and 5). We report correlations for all genotypes and for epistatic genotypes separately.

**Figure 4.**
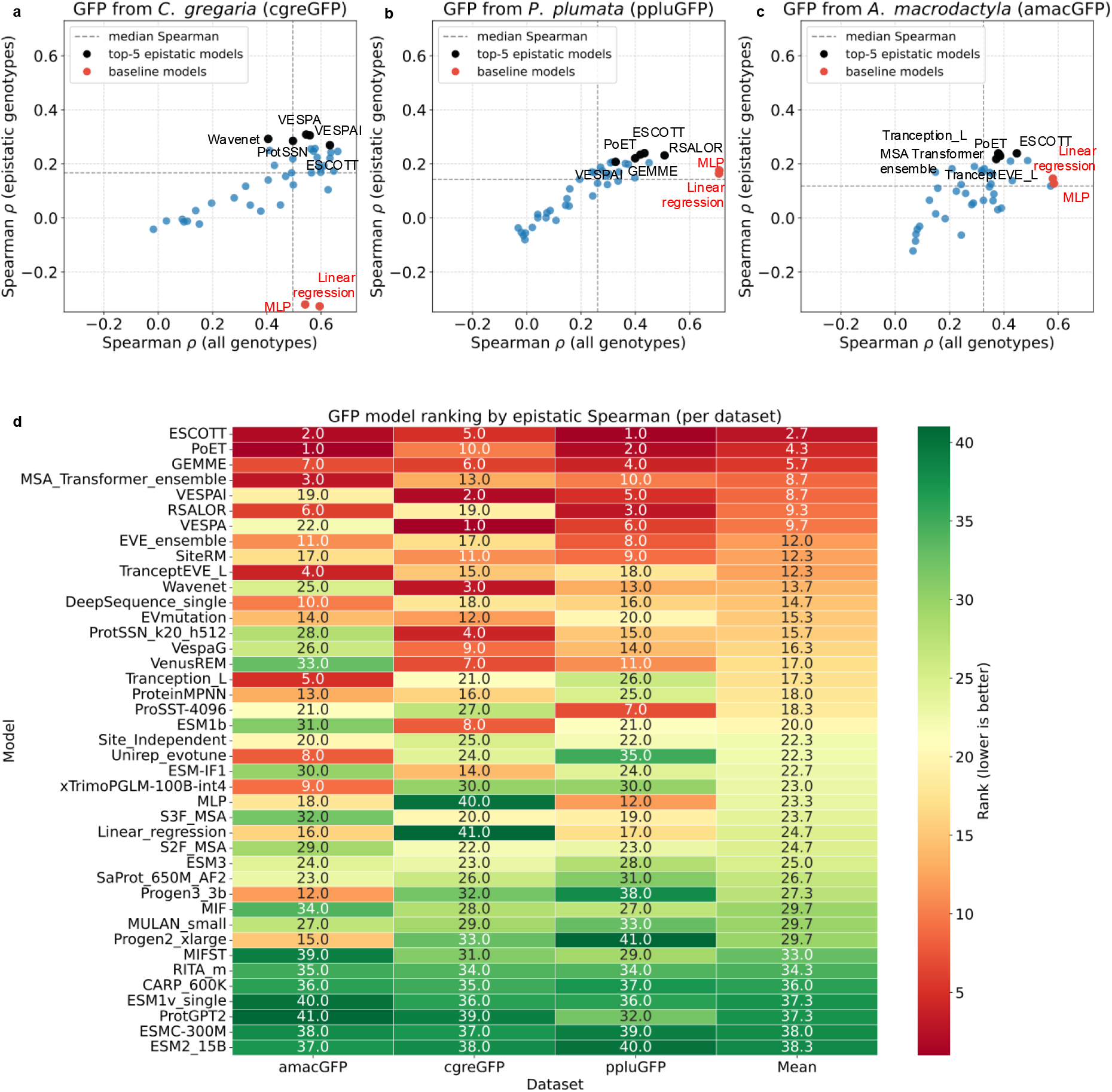
Performance of the zero-shot methods for the three GFP datasets from Somermeyer et al.^11^. The agreement with experiment is measured by Spearman’s rank correlation (ρ) between model predictions and experimental values. Correlations are computed separately for epistatic genotypes (y axis) and a size-matched random sample of all multi-mutant genotypes (x axis). The baseline models are shown as red dots, top five performing models as black dots with labels, all other models are represented as blue dots. Within each model family (e.g., ESM-2 8M, 35M, etc.), the best-performing variant is retained, yielding 39 models per dataset. Baseline linear regression and MLP are shown on the right. **a**. *C. gregaria* (cgreGFP). **b**. *P. plumata* (ppluGFP). **c**. *A*.*macrodactyla* (amacGFP). **d**. Ranking of methods for the GFP datasets. For each method, its performance rank (based on Spearman’s correlation for the epistatic genotypes) was calculated for the three GFP datasets, then the average rank was calculated (last column). Methods are ordered based on the average rank.

**Figure 5.**
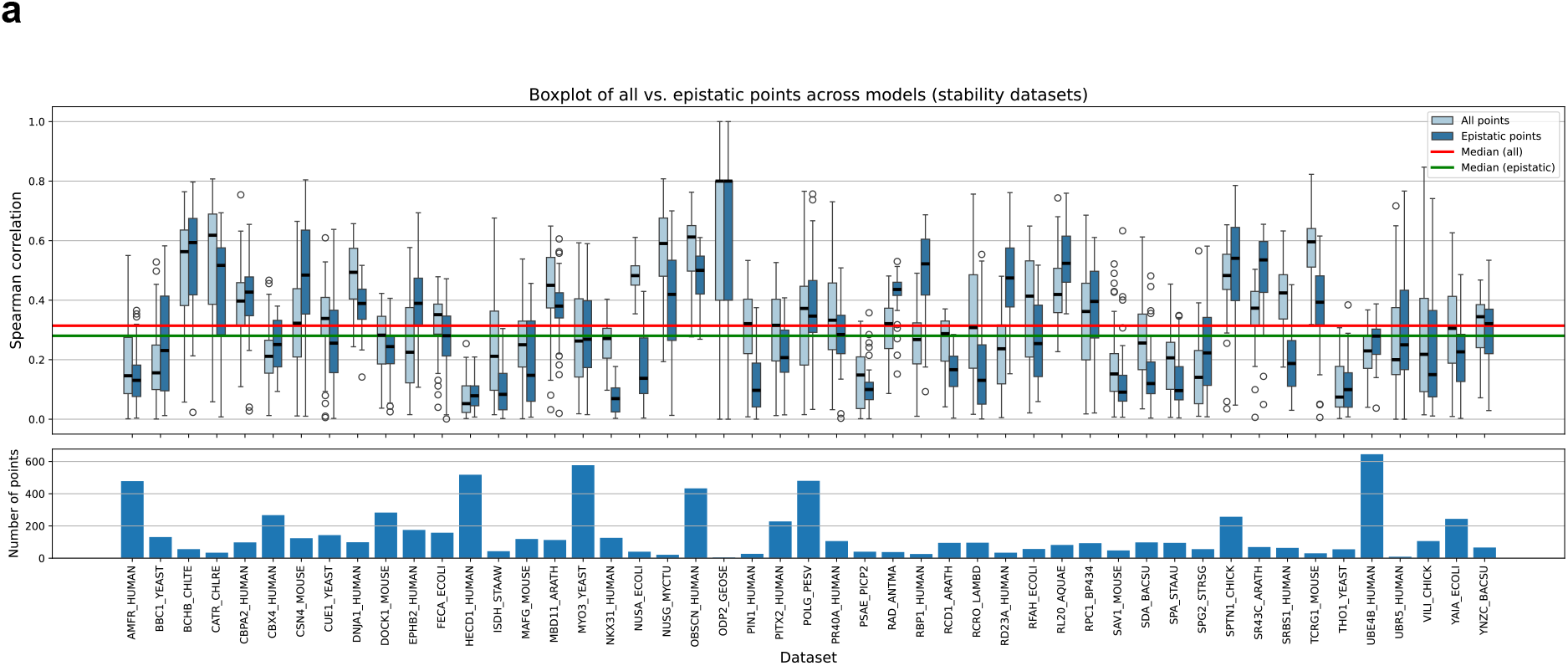
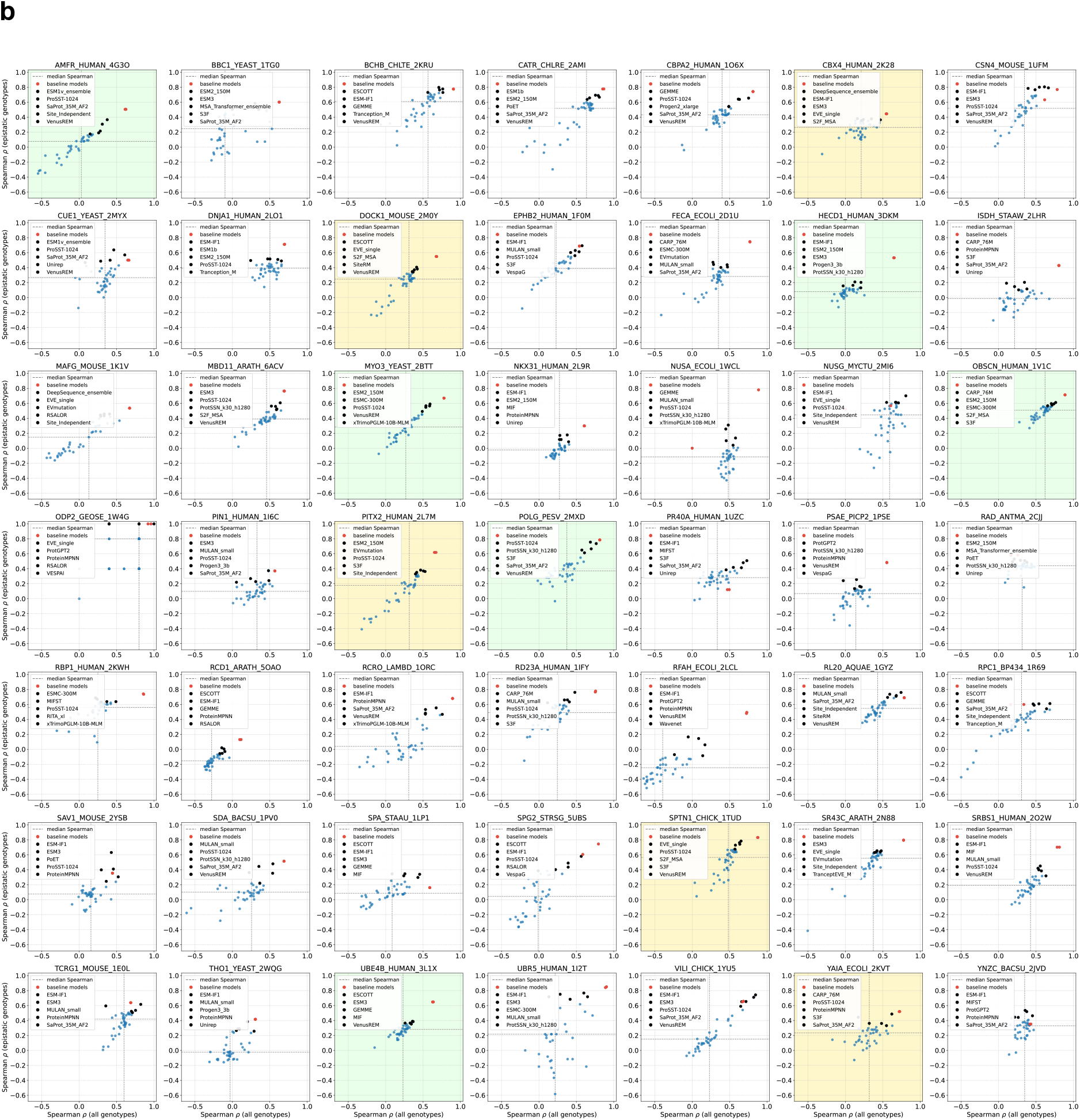
**a**. Performance of zero-shot models for the stability datasets^25^. Correlations are computed separately for epistatic genotypes (dark blue) and a size-matched random sample of all multi-mutant genotypes (light blue). The lower panel shows the number of epistatic genotypes for each dataset. **b**. Spearman’s correlation with experimental data for all genotypes (x axis) and epistatic genotypes (y axis) for the stability datasets. The baseline models (linear regression and MLP) are shown as red dots, the top five performing modes for epistatic genotypes as black dots, all other models are represented as blue dots. Datasets with more than 400 epistatic genotypes are shown on a green background, datasets with the number of epistatic genotypes between 200 and 400 on a yellow background.

#### Somermeyer GFP datasets

For the three GFP datasets, we observe a reasonable agreement of predicted and experimentally measured brightness values for the ‘all genotypes’ set, with the best methods reaching the Spearman’s correlation over 0.6 between predicted and experimental values. At the same time, there is an extremely poor agreement for the epistatic genotypes, where any method rarely exceeds a correlation value of 0.2 (Figure 4a-c, Supplementary Table 2).

For comparison, we trained two simple baseline supervised models using only data on fitness effects of single amino acid mutations: a linear regression model and a multi-layer perceptron (MLP) (red dots in Figure 4). This is not an entirely fair comparison, since these models have access to the labels for the single amino acid mutants, whereas zero-shot models have no access to any labels. Still, we believe this is a useful baseline. The linear regression model by definition cannot take epistatic interactions between mutations into account, and the MLP model is unlikely to do so, since it has been trained on single mutants. So, performance of these two models should be regarded as the minimal baseline, and only models exceeding this baseline can be seen as capable of predicting epistatic effects. Unexpectedly, both models perform comparable or better than zero-shot models for the all genotypes set, again with this effect larger for *P. plumata*. For epistatic genotypes, the performance is comparable to zero-shot models for *A. macrodactyla* and *P. plumata*, but they fail completely for the *C. gregaria* dataset. Hence, we conclude that, first, as a rule, zero-shot VEP models cannot predict epistatic interactions of variants for the fluorescence phenotype, and second, the *C. gregaria* dataset is highly epistatic where linear models cannot perform at all, in accordance with an earlier observation by Somermeyer et al.^11^.

Our data indicate that only a handful of models are able to improve over the baseline, and none perform consistently well across all three datasets. Among them, ESCOTT^26^ and PoET^27^ are among the top five in two of the three datasets and top performers overall (Figure 4d). Both these models make use of homologs of the query sequence, stressing the role of evolutionary conservation for variant effect prediction: ESCOTT does so by considering features derived from amino acid conservation, and PoET works by considering several related sequences in one context window. One other top-performing method, MSA-Transformer^28^, accepts a protein sequence alignment directly as input. Additionally, ESCOTT employs features derived from protein three-dimensional structure, signifying that this is also a valuable source of information for predicting variant effects. Moreover, ESCOTT attempts to take into account epistasis directly by adopting features produced by the GEMME^18^ approach – counting the number of changes required for the query mutation to be accepted by the sequence based on the phylogenetic tree of the protein family – which alone occupies rank three for the GFP datasets. Interestingly, ESCOTT and GEMME are not machine-learning, but purely statistical models. Hence, we argue that clever feature engineering is more important for predicting fitness of epistatic genotypes than complex model architecture. Interestingly, for *C. gregaria*, two classical language models, VESPA and VESPAl^29^ – that are derived by appending an ensemble regression model downstream of several protein language models (PLMs) – performed best, but their performance for epistatic genotypes is poor (Figure 4a). Taken together, these observations allow us to propose that zero-shot methods, specifically PLMs, still lack performance predicting genotype-to-phenotype relationships for epistatic genotypes.

#### Tsuboyama stability datasets

For the stability datasets, the number of epistatic genotypes varies between 4 and 645 in the datasets for different proteins. We focused on datasets containing more than 200 epistatic genotypes based on assessment of thermodynamic coupling (Supplementary Figure 7, see Methods for details). However, when inspecting the thermodynamic coupling distributions in these datasets, we observe that not all of them have a pronounced heavy tail as one would expect for truly epistatic datasets. For example, note the difference between POLG_PESV and SPTN1_CHICK datasets and HECD1_HUMAN and PITX2_HUMAN datasets (Supplementary Figure 8).

Performance of zero-shot methods varies a lot for the different datasets, but in rare cases it exceeds the Spearman’s correlation of 0.6 (Figure 5). To retain statistical power, we again focus on the 11 datasets that contain more than 200 epistatic genotypes (Figure 5b). In them, performance of all models on epistatic genotypes is poor, with no models outperforming the baselines, indicating that epistasis is weak in these datasets. Among datasets with more than 400 epistatic genotypes, for the VPg porcine saprovirus protein involved in viral genome replication (POLG_PESV), the zero-shot models’ performance for epistatic genotypes is just as good as for all genotypes, for some models almost reaching a correlation of 0.8, on par with the baselines. For some datasets, in particular HECD1_HUMAN, a E3 ubiquitin-protein ligase, the performance for all genotypes, as well as for epistatic genotypes is very poor (Spearman’s *ρ* below 0.25), while baseline models perform comparably with other datasets with correlations around 0.6. For the two other E3 ubiquitin-protein ligases, AMFR_HUMAN and UBE4B_HUMAN, all models perform only slightly better with top correlations around 0.4 both for all and for epistatic genotypes. For OBSCN_HUMAN, a muscle structural protein with a kinase activity, there is also a strong correlation between the performances on all and epistatic genotypes, but the baselines here clearly outperform all zero-shot models. For the myosin MYO3_YEAST, some models achieve a relatively good performance of Spearman’s *ρ* 0.6 both for all and epistatic genotypes, but baselines here, again, are better than all zero-shot methods. Among datasets comprising between 200 and 400 epistatic genotypes, only for the structural protein spectrin (SPTN1_CHICK) the median models’ performance on epistatic genotypes is around 0.5 (Figure 5b). Even for the top performing models, the performance for epistatic genotypes is worse than for all genotypes, and much worse than for baseline linear models.

Comparably well performing models for the stability datasets include ProSST^30^, ESM-IF1^31^, VenusREM^32^, SaProt^33^, and S3F^34^ (Figure 6). Just as in the case of GFP datasets, successful models need additional information, such as protein three-dimensional structure (ProSST, VenusREM, SaProt, S3F). Some models (ProSST and its successor VenusREM) additionally consider evolutionary information by including an alignment modality. The other model in the top five is ESM-IF1, which is an inverse folding model: it predicts sequences given a three-dimensional structure (in VEP tasks, an AlphaFold-predicted structure is used^35^), and thus also relies on structural information about the protein. In concordance with this, ESM-IF1 compares favourably to pure sequence-based zero-shot methods for VEP stability tasks where all genotypes are considered disregarding epistasis^36^. Hence one can conclude that three-dimensional structure is essential, but not sufficient (given the overall poor performance of all models) for predicting stability of epistatic genotypes. Surprisingly, there is no overlap between top-performing methods for the GFP and the stability datasets, underscoring the fundamental difference between these phenotypes.

**Figure 6.**
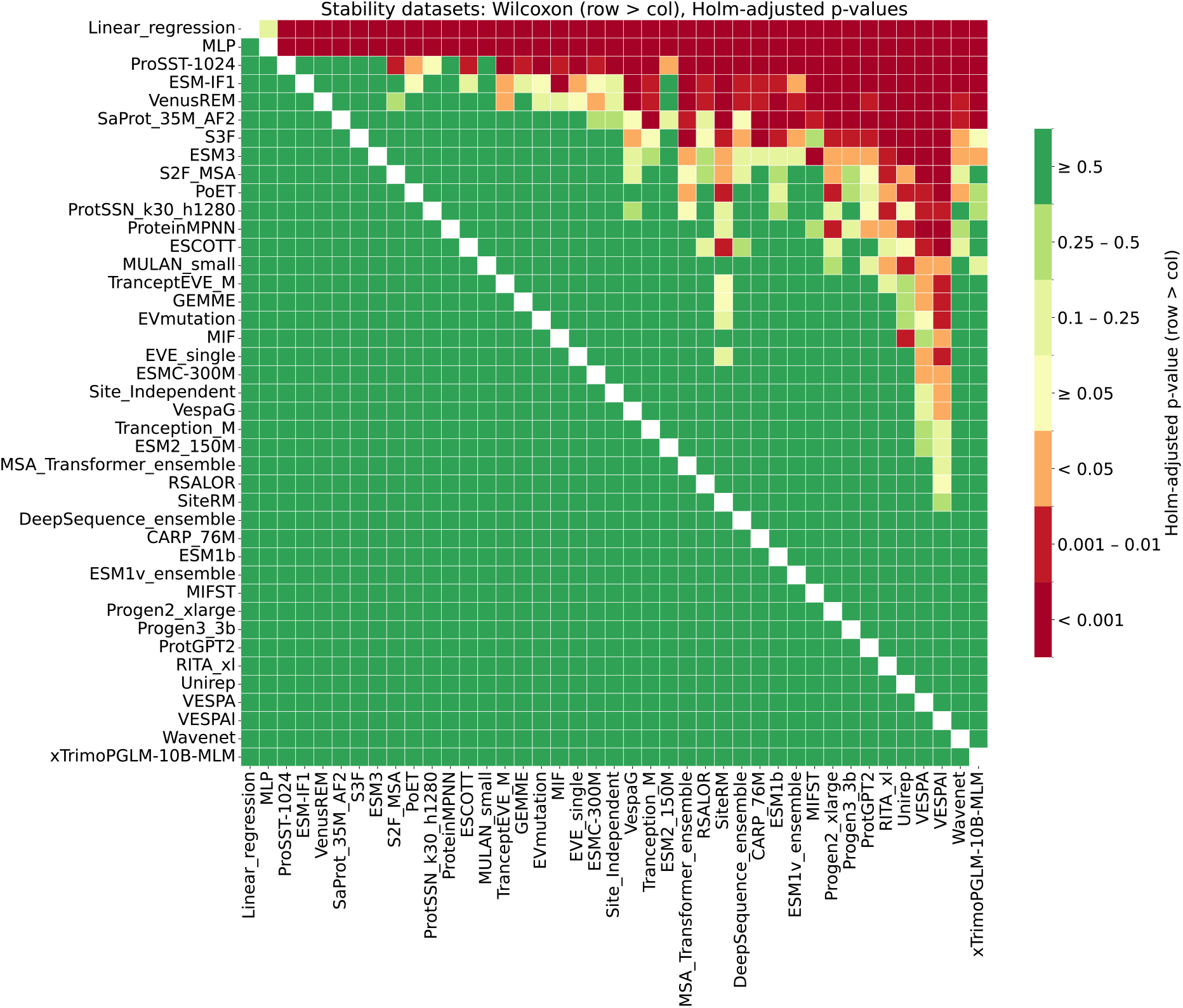
Comparison of zero-shot methods for the Tsuboyama stability datasets. Every pair of methods was compared based on their performance for all datasets (see Methods for details). Methods are sorted from best to worst (based on the sum of the values in a row from lowest to highest, alphabetically in case of a tie) with the statistical significance color-coded from red (highly significant) to green (non-significant).

## Discussion

Epistatic interactions between mutations in a protein sequence is an ubiquitous biological phenomenon, and understanding and predicting their effect correctly is a major challenge in computational and evolutionary biology. MAVE experiments provided an unprecedented amount of data for training computational variant effect prediction (VEP) tools. Despite these data being large, epistasis is still not well captured in them. Indeed, of 217 MAVE datasets from the ProteinGym benchmark^23^, only 69 contain data on multi-mutational variants, of which only GFP datasets contain a considerable number of variants with more than two mutations. Interestingly, even this limited amount of data led to creation of several AI models capable of designing new functional GFP variants^37–39^. Nevertheless, to date no model can truly traverse the sequence landscape and, for example, find functional proteins on a path connecting one species to another^11^.

In this work, we assessed the capability of state-of-the-are VEP methods to predict effects of epistatic interactions correctly and focused our effort on the 52 MAVE datasets where we could confidently estimate the extent of epistasis from the experimental data including experimental errors. A striking conclusion of our study is that virtually none of the existing VEP tools are able to predict epistatic effects out of the box. While these models perform adequately on datasets containing single mutations and mutations whose effects combine linearly, their performance declines significantly when tasked with predicting sequences containing epistatic combinations of mutations. Several models show medium-to-high success rates in isolated cases, and these are usually top-performing models on the whole ProteinGym dataset (for example, at the time of this writing these include VenusREM^40^, ProSST^41^, ESCOTT^26^). Interestingly, these models typically consider features derived from protein structure and evolution (in some cases direct epistatic terms from GEMME^18^) but not necessarily have a complex architecture: for example, ESCOTT is a statistical model with no machine learning in it. This highlights that a prudent treatment of data is the key to successful VEP prediction, and not complex deep learning.

Importantly, many zero-shot models – protein language models (PLMs) in the first place – are not specifically trained for the VEP task. A classical PLM is trained to predict masked tokens in a protein sequence based on information from all other (unrelated) proteins in the Universe. They can be re-purposed as a VEP tool in that they predict how likely a mutated protein sequence is given all that it knows about naturally occurring proteins. That is, a PLM predicts some abstract notion of plausibility of a sequence instead of assessing the particular set of mutations. Our results here suggest that this notion is close enough to the real functional effect for single mutants and simple linear-effect combinations of single mutants, yet it fails for more complex interactions between variants, such as epistasis. However, many current VEP models still rely on PLMs at least as a starting point. Recent reports suggest that it is necessary to include non-linear transformations of the data to achieve some success in predicting epistasis^42^. Unfortunately, to date these studies are anecdotal and confined to a handful of proteins, not allowing to paint a broad picture across a variety of proteins with various functions.

Hence, we demonstrated that the notion of sequence evolutionary plausibility learned by zero-shot models does not generalize well to distant regions in the sequence space, and in particular cannot cross low-fitness valleys (Figure 1a). One explanation to this can be that PLMs learn pairwise evolutionary interactions between position^43^, and epistatic effects are of a higher dimension. Another reason could be that zero-shot models have only seen evolutionary competent (functional) protein sequences, which occupy a tiny fraction of the sequence space, and thus have no information about vast regions of it. In another work^44^, we demonstrated that zero-shot methods rather capture a phenotype related to mutations’ pathogenicity than direct functional impact on protein activity. Taken together, all this may contribute to the failure of such methods to predict the phenotype for highly epistatic genotypes.

Accurate prediction of epistatic effects is crucial for many practical applications, such as protein design, as well as for fundamental understanding of protein evolution. Our results suggest that further refinement of current models or development of new architectures specifically tailored for multi-mutational and epistatic data is necessary. While current zero-shot models show promise in predicting the effects of individual mutations, our study highlights the need for more specialized approaches to accurately capture epistasis. Addressing this gap is critical for advancing both theoretical understanding and practical applications in protein science.

## Methods

### Preprocessing

#### Dataset of Somermeyer et al

For each genotype, we first estimated experimental error by calculating the weighted mean and weighted standard deviation using the nucleotide genotypes given the corresponding values of pseudocell_count (total_cell_count), replicates_mean_brightness (brightness) and replicates_stdev_weighted (brightness_stdev) from the original data files. We computed the resulting brightness standard deviation brightness_stdev for a certain amino acid sequence of GFP, which we call an amino acid genotype, as 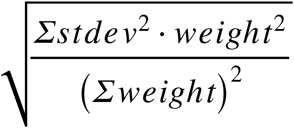, where *stdev* and *weight* are corresponding brightness_stdev and total_cell_count, for each of the nucleotide genotypes corresponding to a certain protein sequence. For the brightness mean of an amino acid genotype, we computed a weighted mean of brightnesses of nucleotide genotypes. We then scaled the resulting values against the wild type, so we subtracted the mean brightness of WT from the brightness, and for the brightness standard deviation, we 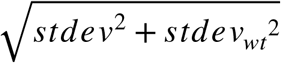 took *stdev*^2^ + *stdev*_*wt*_^2^. The wildtype sequences were then removed from the data.

#### Dataset of Tsuboyama et al

Firstly, we removed datasets containing only single mutants, which yielded 50 datasets with double mutants present in the data. We obtained deltaG, deltaG_95CI_low and deltaG_95CI_high for each genotype from Tsuboyama2023_Dataset2_Dataset3_20230416.csv, where deltaG is the median of the posterior probability of *ΔG*, deltaG_95CI_low is the top 2.5%ile posterior of *ΔG* and deltaG_95CI_high is the top 97.5%ile posterior of *ΔG* , which correspond to the median, lower and higher boundary of the credible interval (CI). The length of the left part of CI was computed as left = median - low and right part as right = high - median. We checked if the interval is symmetric: min(left, right) /max(left, right) >= 0.9. If it was, we derived the standard deviation as (high - low) /(2 * 1.96), in other case as (left + right) /(2 * 1.96).

We considered deltaG as observed effects, while the expected effects were reconstructed deltaG, which we obtained by running the thermodynamic model provided in the Supplementary Information of Tsuboyama et al.^25^ (section Thermodynamic coupling analysis). Thermodynamic coupling was then calculated, which is the difference between deltaG and reconstructed deltaG and was used as epistasis in the corresponding formula.

### Epistasis detection

We used the formula (2) as a general formula to detect epistatic genotypes. The intuition behind it is that the difference between the observed phenotype (fluorescence or thermostability) and the one expected from a linear model should be greater than the sum of estimated experimental error times a factor *N* The choice of *N* is described in Supplementary Note 1.

In case of both datasets, we computed the expected effect as the sum of effects for the corresponding single mutants corresponding to each mutation observed in the multi-mutant genotype. We considered only amino acid genotypes throughout this study. If any required information for one of the single mutants was missing, we discarded this genotype. If the expected effect was below the minimum observed effect, we took the latter instead.

### Model performance evaluation

To evaluate zero-shot methods for VEP, we merged the genotypes with the corresponding datasets from zero_shot_substitutions_scores from ProteinGym^23^. It is worth mentioning that sequence numbering in the Somermeyer datasets should be shifted by one position in order to match with the ProteinGym data. We called genotypes epistatic if the epistasis could have been calculated (no missing single mutant genotypes data) and if it passed the criterion described by the epistasis formula (2). Next, for each model, we computed Spearman correlation between the DMS_score (true value) and the model predictions for all epistatic genotypes. We then sampled randomly as many sequences from all multi-mutant genotypes as there were epistatic genotypes, and calculated Spearman correlation for this subset of genotypes. Seed for randomization (pandas.DataFrame.sample) was fixed as 42 except for the whiskers barplots, where we evaluated on five random seeds. As the difference was not significant, we proceeded with one seed strategy for all further computations. As ProteinGym contains predictions for several versions there are many parameter combinations for several models (e.g., the ESM2 family includes ESM2_8M, ESM2_35M, etc.), we selected the best variant for each model from the constructed table with Spearman correlation scores. To do so, we first filled the missing values with 0. Next, for each model variant, the mean of “all genotypes”-values was taken, and the variant with the highest mean was considered best.

The dataset TNKS2_HUMAN was removed from the analysis, as it contained only one epistatic sequence.

### Baselines

#### Linear regression

We used the default linear regression model from scikit-learn and trained it using all single-mutation genotypes, leveraging one-hot encoding for the amino acid sequences. In addition, for the GFP datasets, we log-transformed the target value for training due to its scale, and reversed it for predictions. Then, we computed Spearman correlations as described above. Unless explicitly stated otherwise, we sampled randomly from all multi-mutant genotypes the same number of sequences as there were epistatic genotypes five times, and as final correlation provided the mean across the five seeds.

#### Multilayer perceptron

We used a simple MLP model consisting of two fully connected hidden layers with 10 and 100 neurons, respectively, each followed by a ReLu activation function. The output layer is a single linear unit. We used the Adam optimizer with a learning rate of 1e-3. The loss was MSE and the batch size was 64. The model was trained on 500 epochs and was terminated early if the training loss did not improve for 10 consecutive epochs (early stopping with patience = 10). Pytorch^45^ was used for implementation. For calculating the resulting Spearman correlations, we used the same strategy as for linear regression, fixing the random seed at 42.

#### Statistical evaluation of models on stability datasets

We compared models using the columns of Spearman correlations on epistatic subsets of Tsuboyama stability dataset. Across datasets, we first ran a Friedman test (friedmanchisquare in the scipy.stats package) to assess overall differences among models, as suggested by^46^. When the omnibus test was significant (p-value 3.28e-95), we performed pairwise one-sided Wilcoxon signed-rank tests (as implemented in scipy.stats) on per-dataset *ρ* differences with the alternative *H*_1_: *model*_*i*_ > *model*_*j*_ (median difference > 0); exact zero differences were excluded (zero_method=‘wilcox’). To control the family-wise error rate over all directed comparisons, p-values were adjusted by the Holm step-down procedure. We visualized the Holm-adjusted p-values in a models×models heatmap; the diagonal was left blank, and rows/columns were ordered by the row-wise sum of adjusted p-values (lower sum nearer the top, ties broken alphabetically). Colors were shown in discrete bins with a pivot at *α=0*.*05* (*ρ* < *α* in red hues; *ρ* ≥ *α* in green hues).

## Data availability

Model scores, sequences and labels were obtained from ProteinGym (https://proteingym.org/) in June 2025 via link https://marks.hms.harvard.edu/proteingym/ProteinGym_v1.3/zero_shot_substitutions_scores.zip. For additional experimental errors of GFP, we used https://github.com/aequorea238/Orthologous_GFP_Fitness_Peaks/blob/master/data/final_datasets/amacGFP_cgreGFP_ppluGFP2_final_nucleotide_genotypes_to_brightness.csv from the Somermeyer paper, as well as WT protein sequences from https://github.com/aequorea238/Orthologous_GFP_Fitness_Peaks/blob/master/data/alignments/vGFP_amacGFP_cgreGFP_ppluGFP2__protein_sequences.fasta. For simplicity, we translated the full GFP names into short forms everywhere. For extracting the dG values in the Tsuboya mast ability dataset ,were ferred to the file Tsuboyama2023_Dataset2_Dataset3_20230416.csv inside Processed_K50_dG_datasets.zip from https://doi.org/10.5281/zenodo.7992926 from the original paper.

## Code availability

The code for reproducing results is available on Github: https://github.com/kalininalab/epistasis_proteingym.

## Acknowledgements

We are grateful to Dr. Olga O. Bochkareva, Dr. Carene Antonella Benasolo, and Dr. Alexander Gress for a critical reading of the manuscript.

## References

1. Chaurasia, S. & Dutheil, J. Y. The Structural Determinants of Intra-Protein Compensatory Substitutions. Mol. Biol. Evol. 39, msac063 (2022).

2. Sackton, T. B. & Hartl, D. L. Genotypic Context and Epistasis in Individuals and Populations. Cell 166, 279–287 (2016).

3. Ivankov, D. N., Finkelstein, A. V. & Kondrashov, F. A. A structural perspective of compensatory evolution. Curr. Opin. Struct. Biol. 26, 104–112 (2014).

4. Kondrashov, A. S., Sunyaev, S. & Kondrashov, F. A. Dobzhansky-Muller incompatibilities in protein evolution. Proc. Natl. Acad. Sci. U. S. A. 99, 14878–14883 (2002).

5. Sarkisyan, K. S. et al. Local fitness landscape of the green fluorescent protein. Nature 533, 397–401 (2016).

6. Soskine, M. & Tawfik, D. S. Mutational effects and the evolution of new protein functions. Nat. Rev. Genet. 11, 572–582 (2010).

7. Bloom, J. D., Gong, L. I. & Baltimore, D. Permissive Secondary Mutations Enable the Evolution of Influenza Oseltamivir Resistance. Science 328, 1272–1275 (2010).

8. Papkou, A., Garcia-Pastor, L., Escudero, J. A. & Wagner, A. A rugged yet easily navigable fitness landscape. Science 382, eadh3860 (2023).

9. Pokusaeva, V. O. et al. An experimental assay of the interactions of amino acids from orthologous sequences shaping a complex fitness landscape. PLOS Genet. 15, e1008079 (2019).

10. Vaishnav, E. D. et al. The evolution, evolvability and engineering of gene regulatory DNA. Nature 603, 455–463 (2022).

11. Gonzalez Somermeyer, L. et al. Heterogeneity of the GFP fitness landscape and data-driven protein design. eLife 11, e75842 (2022).

12. Poelwijk, F. J., Tănase-Nicola, S., Kiviet, D. J. & Tans, S. J. Reciprocal sign epistasis is a necessary condition for multi-peaked fitness landscapes. J. Theor. Biol. 272, 141–144 (2011).

13. DePristo, M. A., Weinreich, D. M. & Hartl, D. L. Missense meanderings in sequence space: a biophysical view of protein evolution. Nat. Rev. Genet. 6, 678–687 (2005).

14. Fröhlich, C. et al. Epistasis arises from shifting the rate-limiting step during enzyme evolution of a β-lactamase. Nat. Catal. 7, 499–509 (2024).

15. Gerasimavicius, L., Teichmann, S. A. & Marsh, J. A. Leveraging protein structural information to improve variant effect prediction. Curr. Opin. Struct. Biol. 92, 103023 (2025).

16. Bromberg, Y., Prabakaran, R., Kabir, A. & Shehu, A. Variant Effect Prediction in the Age of Machine Learning. Cold Spring Harb. Perspect. Biol. a041467 (2024) doi:10.1101/cshperspect.a041467.

17. Meier, J. et al. Language models enable zero-shot prediction of the effects of mutations on protein function. 2021.07.09.450648 Preprint at 10.1101/2021.07.09.450648 (2021).

18. Laine, E., Karami, Y. & Carbone, A. GEMME: A Simple and Fast Global Epistatic Model Predicting Mutational Effects. Mol. Biol. Evol. 36, 2604–2619 (2019).

19. Cheng, J. et al. Accurate proteome-wide missense variant effect prediction with AlphaMissense. Science 381, eadg7492 (2023).

20. Rives, A. et al. Biological structure and function emerge from scaling unsupervised learning to 250 million protein sequences. Proc. Natl. Acad. Sci. 118, e2016239118 (2021).

21. The UniProt Consortium. UniProt: the Universal Protein Knowledgebase in 2025. Nucleic Acids Res. 53, D609–D617 (2025).

22. McEwen, A. E., Tejura, M., Fayer, S., Starita, L. M. & Fowler, D. M. Multiplexed assays of variant effect for clinical variant interpretation. Nat. Rev. Genet. 27, 137–154 (2026).

23. Notin, P. et al. ProteinGym: Large-Scale Benchmarks for Protein Fitness Prediction and Design. Adv. Neural Inf. Process. Syst. 36, 64331–64379 (2023).

24. Wright, S. The roles of mutation, inbreeding, crossbreeding and selection in evolution. Proc. Sixth Int. Congr. Genet. 1, 356–366 (1932).

25. Tsuboyama, K. et al. Mega-scale experimental analysis of protein folding stability in biology and design. Nature 620, 434–444 (2023).

26. Tekpinar, M., David, L., Henry, T. & Carbone, A. PRESCOTT: a population aware, epistatic, and structural model accurately predicts missense effects. Genome Biol. 26, 113 (2025).

27. Truong Jr, T. & Bepler, T. PoET: A generative model of protein families as sequences-of-sequences. Adv. Neural Inf. Process. Syst. 36, 77379–77415 (2023).

28. Rao, R. M. et al. MSA Transformer. in Proceedings of the 38th International Conference on Machine Learning 8844–8856 (PMLR, 2021).

29. Marquet, C. et al. Embeddings from protein language models predict conservation and variant effects. Hum. Genet. 141, 1629–1647 (2022).

30. Li, M. et al. ProSST: Protein Language Modeling with Quantized Structure and Disentangled Attention. Adv. Neural Inf. Process. Syst. 37, 35700–35726 (2024).

31. Hsu, C. et al. Learning inverse folding from millions of predicted structures. 2022.04.10.487779 Preprint at 10.1101/2022.04.10.487779 (2022).

32. Tan, Y., Wang, R., Wu, B., Hong, L. & Zhou, B. From high-throughput evaluation to wet-lab studies: advancing mutation effect prediction with a retrieval-enhanced model. Bioinformatics 41, i401–i409 (2025).

33. Su, J. et al. SaProt: Protein Language Modeling with Structure-aware Vocabulary. 2023.10.01.560349 Preprint at 10.1101/2023.10.01.560349 (2024).

34. Zhang, Z. et al. Multi-Scale Representation Learning for Protein Fitness Prediction. Preprint at 10.48550/arXiv.2412.01108 (2024).

35. Jumper, J. et al. Highly accurate protein structure prediction with AlphaFold. Nature 596, 583–589 (2021).

36. Paul, S., Kollasch, A., Notin, P. & Marks, D. Combining Structure and Sequence for Superior Fitness Prediction. in (2023).

37. Weinstein, J. Y. et al. Designed active-site library reveals thousands of functional GFP variants. Nat. Commun. 14, 2890 (2023).

38. Gelman, S. et al. Biophysics-based protein language models for protein engineering. Nat. Methods 22, 1868–1879 (2025).

39. Bigge, B. M., Lane, R., McGeever, E., Wood, H. & York, R. Efficient GFP variant design with a simple neural network ensemble. The Stacks 10.57844/arcadia-tuvv-w59k (2025) doi:10.57844/arcadia-tuvv-w59k.

40. Tan, Y., Wang, R., Wu, B., Hong, L. & Zhou, B. From high-throughput evaluation to wet-lab studies: advancing mutation effect prediction with a retrieval-enhanced model. Bioinformatics 41, i401–i409 (2025).

41. Li, M. et al. ProSST: Protein Language Modeling with Quantized Structure and Disentangled Attention. 2024.04.15.589672 Preprint at 10.1101/2024.04.15.589672 (2024).

42. Nambiar, A., Littlefield, S. B., Cuellar, C., Khorana, R. & Maslov, S. Protein Language Models Capture Structural and Functional Epistasis in a Zero-Shot Setting. 2025.09.14.676130 Preprint at 10.1101/2025.09.14.676130 (2025).

43. Zhang, Z. et al. Protein language models learn evolutionary statistics of interacting sequence motifs. Proc. Natl. Acad. Sci. 121, e2406285121 (2024).

44. Gress, A. et al. StructGuy: Data leakage free prediction of functional effects of genetic variants. 2025. 12.0 1.691563 Preprint at 10.64898/2025.12.01.691563 (2025).

45. Paszke, A. et al. PyTorch: An Imperative Style, High-Performance Deep Learning Library. Preprint at 10.48550/arXiv.1912.01703 (2019).

46. Demšar, J. Statistical Comparisons of Classifiers over Multiple Data Sets. J Mach Learn Res 7, 1–30 (2006).

