## Supplementary Material for "Beyond additivity: zero-shot methods cannot predict impact of epistasis on protein properties and function"

**a.**

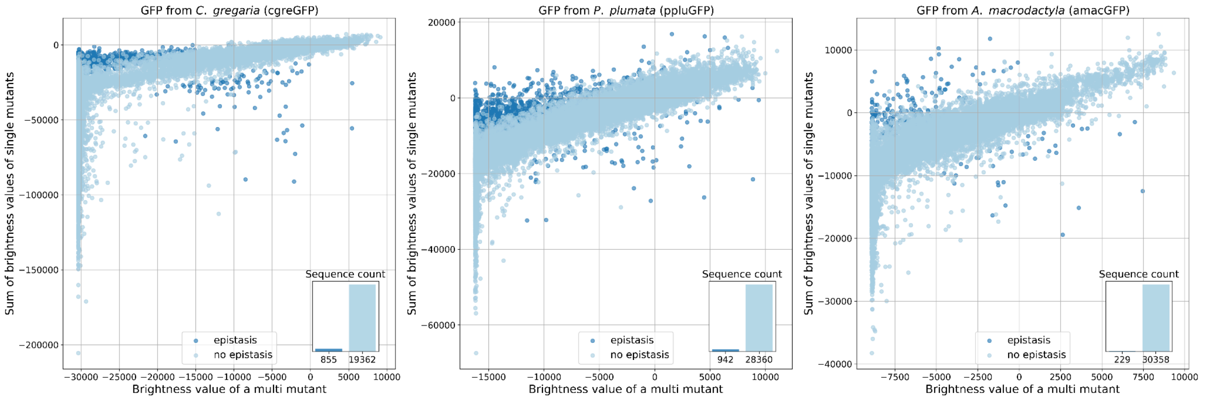


**b.**

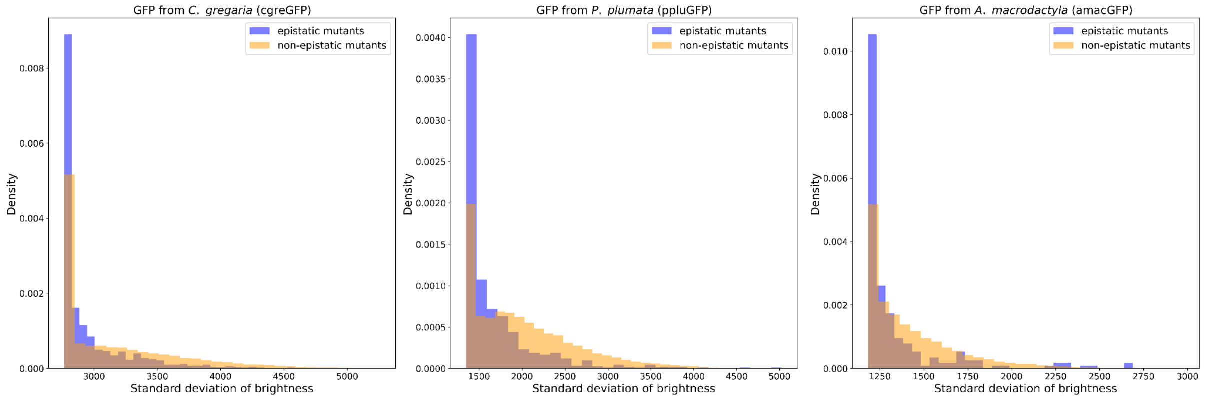


**Supplementary Figure 1. a.** Epistatic and non-epistatic genotypes in the dataset for the three GFP from *C. gregaria*, *P. plumata* and *A. macrodactyla*: for each multi-mutant genotype, the x-axis shows its observed brightness and the y-axis shows the sum of the corresponding single-mutant brightness values. Both values are shown as raw brightness values (not log-transformed) and relative to the wildtype. Dark blue dots correspond to the epistatic genotypes. **b.** The distribution of experimental errors in epistatic (dark blue) and non-epistatic (orange) genotypes for *C. gregaria*, *P. plumata* and *A. macrodactyla*. Observed epistasis cannot be attributed to larger experimental errors, and hence is a genuine biological effect.

**Supplementary Note 1.** Selection of *N*.
We experimented with different values of *N* (see Formula 2 from the main text), separately for the Somermeyer GFP and the Tsuboyama stability datasets. The results for the Somermeyer datasets can be seen in Supplementary Figure 2. In brief, we chose *N* = 1 as the final value as for larger Ns the number of epistatic sequences became too low. For the Tsuboyama datasets (Supplementary Figures 4-5), we considered *N* = 3 to be the optimal value, as for lower *N*s there remained too many epistatic sequences.

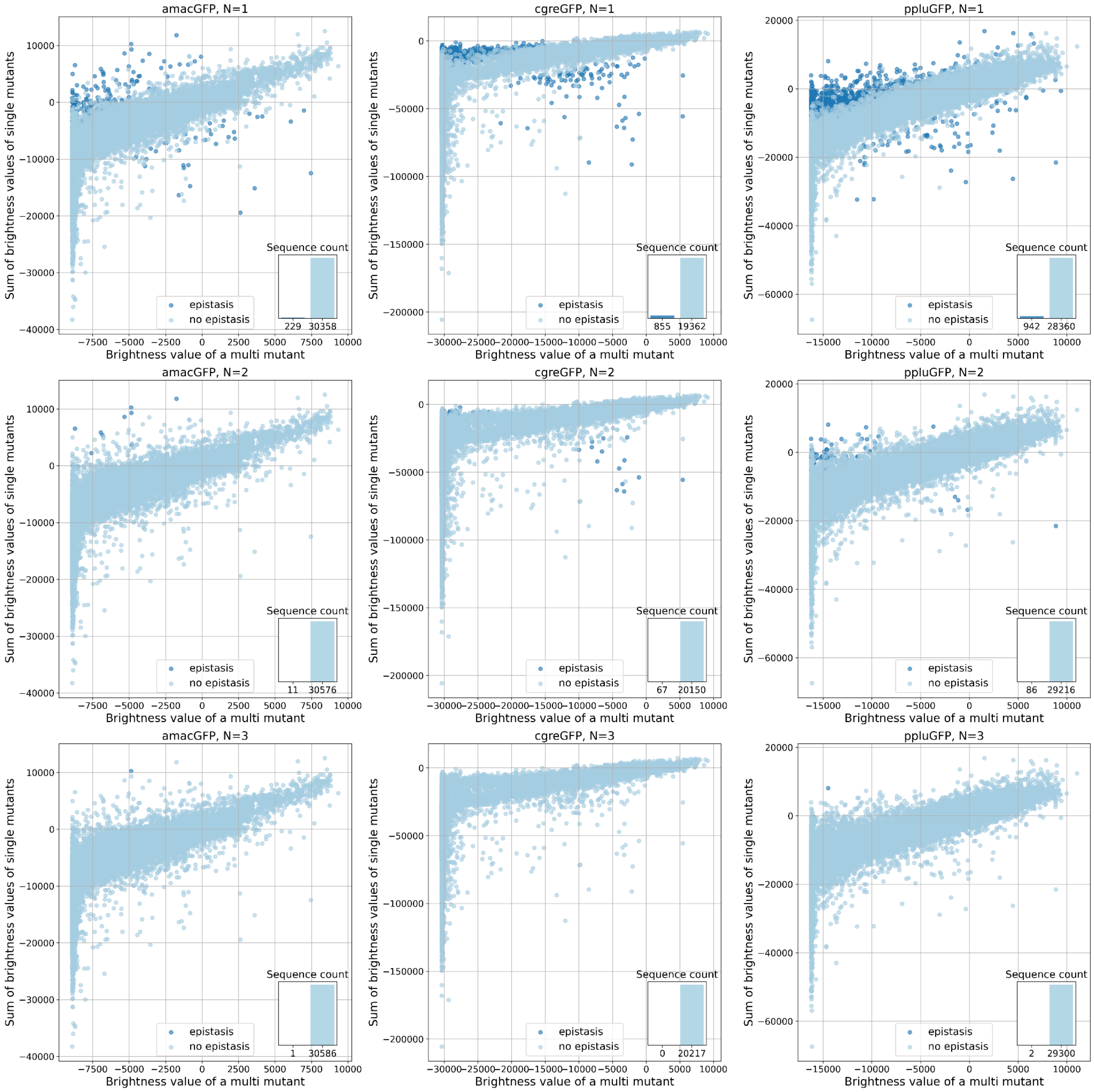


**Supplementary Figure 2.** Epistatic and non-epistatic genotypes in the dataset for the three GFP from *A. macrodactyla*, *C. gregaria* and *P. plumata*, depending on the selection of *N*. Results for N=1, 2, 3 are demonstrated, as well as the sequence count: epistatic and non-epistatic. For each multi-mutant genotype, the x-axis shows its observed brightness and the y-axis shows the sum of the corresponding single-mutant brightness values. Both values are shown as raw brightness values (not log-transformed) and relative to the wildtype. Dark blue dots correspond to the epistatic genotypes, i.e. where the functional effect of a combination of mutations significantly (larger than the experimental error) differs from the linear combination of individual effects.

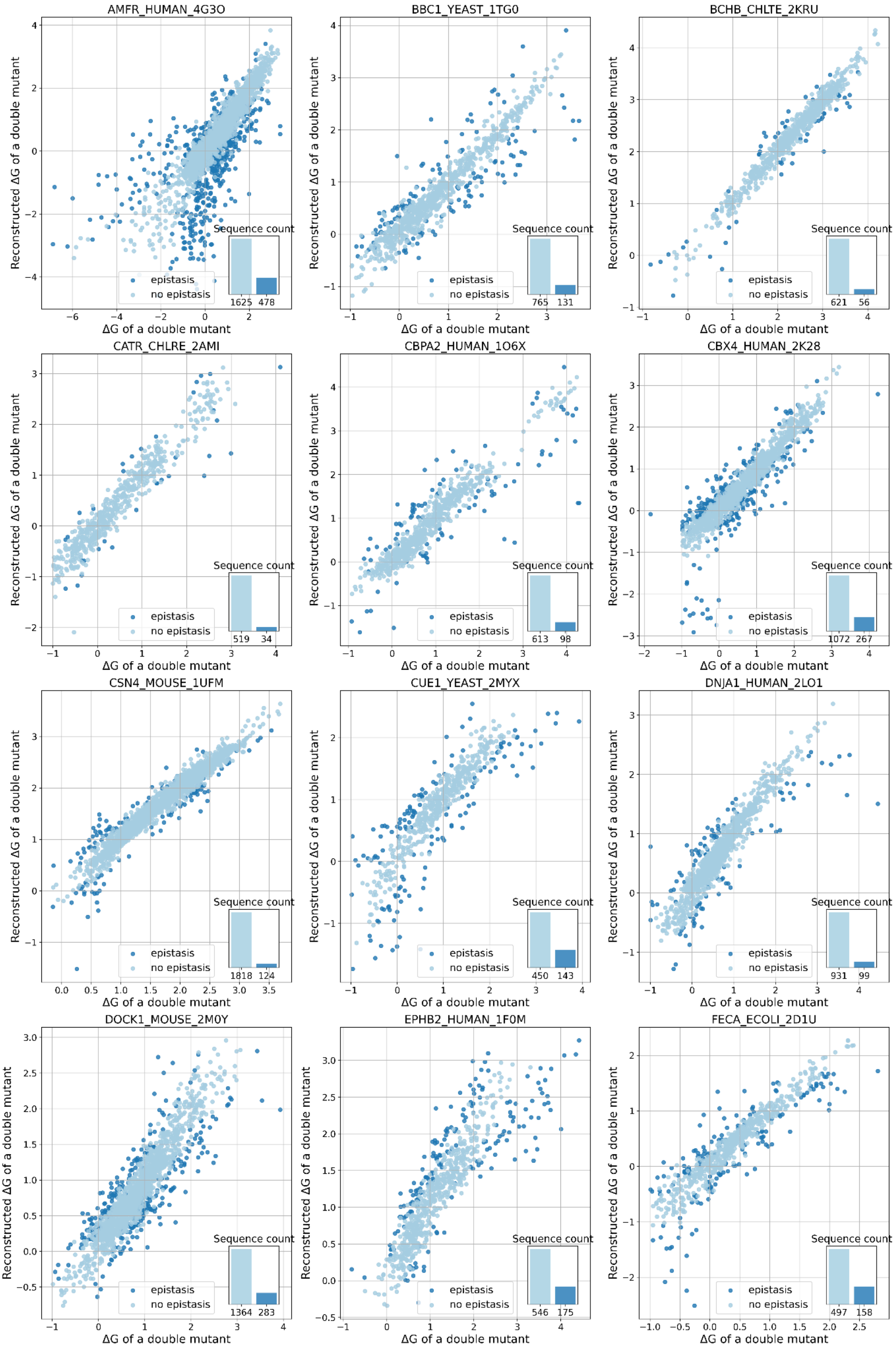
AMFR_HUMAN_4G3O
BBC1_YEAST_1TG0
BCHB_CHLTE_2KRU
CATR_CHLRE_2AMI
CBPA2_HUMAN_1O6X
CBX4_HUMAN_2K28
CSN4_MOUSE_1UFM
CUE1_YEAST_2MYX
DNJA1_HUMAN_2LO1
DOCK1_MOUSE_2M0Y
EPHB2_HUMAN_1F0M
FECA_ECOLI_2D1U
Reconstructed ΔG of a double mutant
ΔG of a double mutant
epistasis
no epistasis
Sequence count
1625 478
765 131
621 56
519 34
613 98
1072 267
1818 124
450 143
931 99
1364 283
546 175
497 158

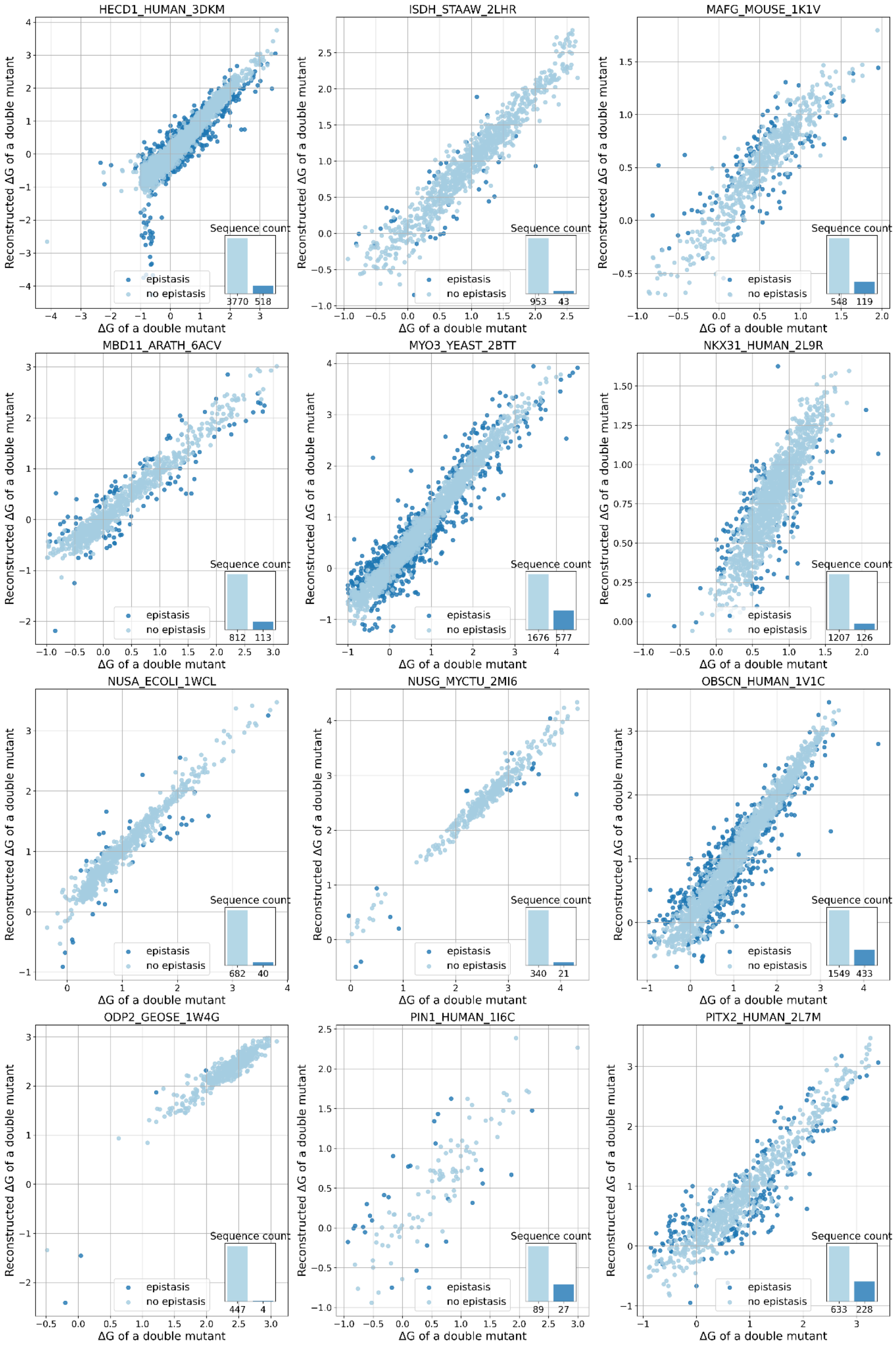

HECD1_HUMAN_3DKM
3770 518
ISDH_STAAW_2LHR
953 43
MAFG_MOUSE_1K1V
548 119
MBD11_ARATH_6ACV
812 113
MYO3_YEAST_2BTT
1676 577
NKX31_HUMAN_2L9R
1207 126
NUSA_ECOLI_1WCL
682 40
NUSG_MYCTU_2MI6
340 21
OBSCN_HUMAN_1V1C
1549 433
ODP2_GEOSE_1W4G
447 4
PIN1_HUMAN_1I6C
89 27
PITX2_HUMAN_2L7M
633 228
Sequence count
epistasis
no epistasis
Reconstructed ΔG of a double mutant
ΔG of a double mutant

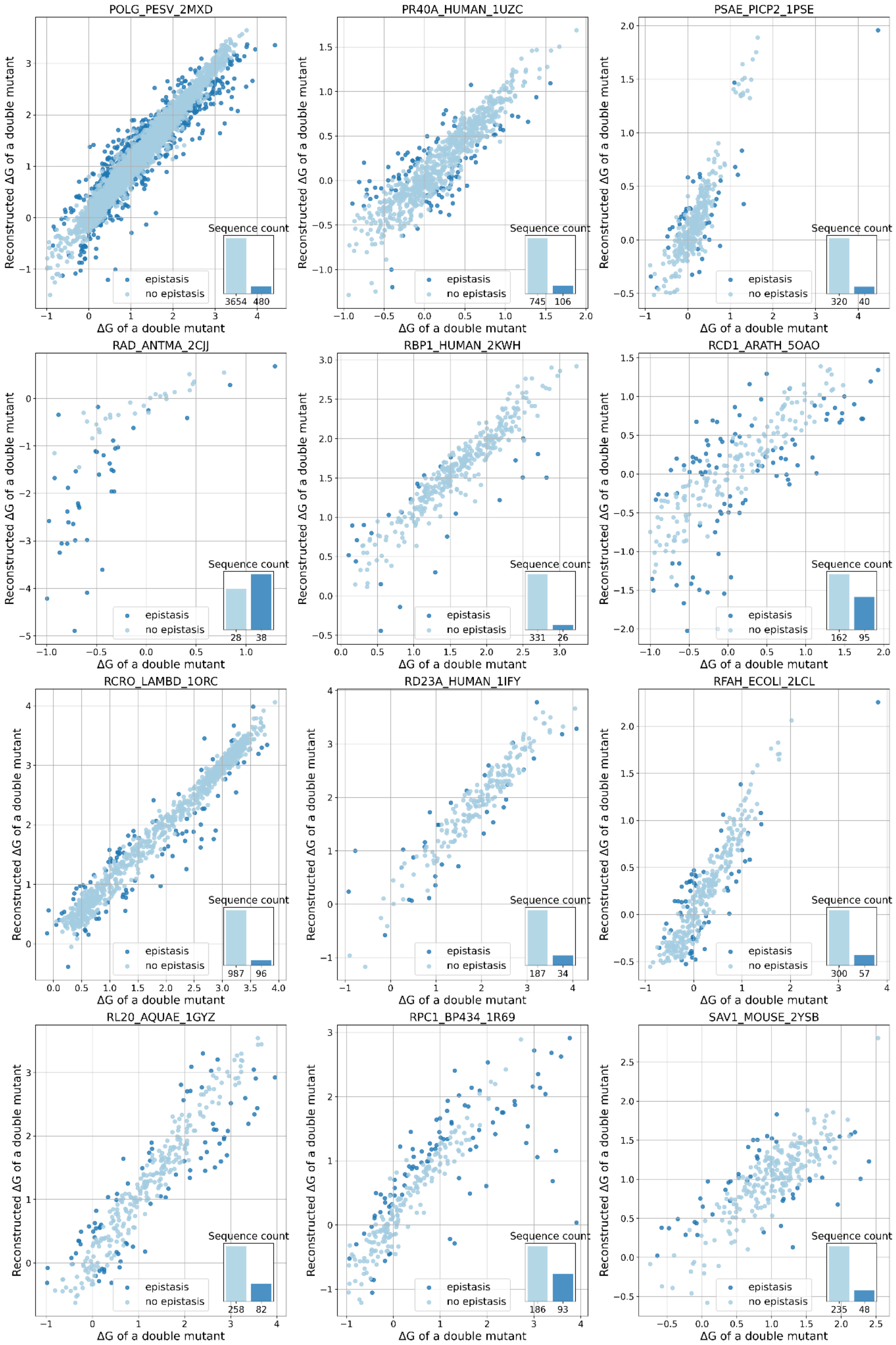

POLG_PESV_2MXD
Reconstructed ΔG of a double mutant
ΔG of a double mutant
Sequence count
epistasis
no epistasis
3654
480
PR40A_HUMAN_1UZC
745
106
PSAE_PICP2_1PSE
320
40
RAD_ANTMA_2CJJ
28
38
RBP1_HUMAN_2KWH
331
26
RCD1_ARATH_5OAO
162
95
RCRO_LAMBD_1ORC
987
96
RD23A_HUMAN_1IFY
187
34
RFAH_ECOLI_2LCL
300
57
RL20_AQUAE_1GYZ
258
82
RPC1_BP434_1R69
186
93
SAV1_MOUSE_2YSB
235
48

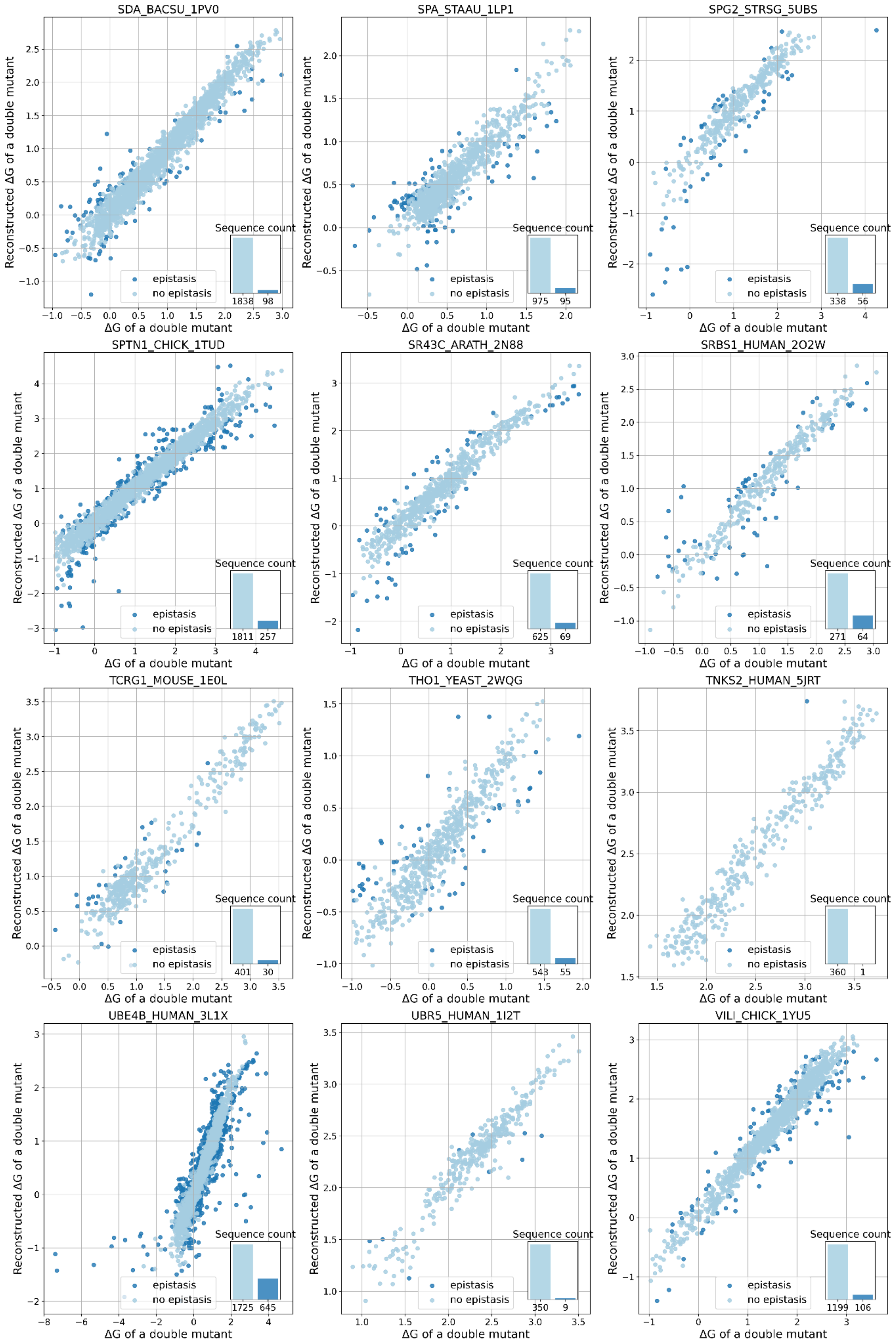
SDA_BACSU_1PV0
SPA_STAAU_1LP1
SPG2_STRSG_5UBS
SPTN1_CHICK_1TUD
SR43C_ARATH_2N88
SRBS1_HUMAN_2O2W
TCRG1_MOUSE_1E0L
THO1_YEAST_2WQG
TNKS2_HUMAN_5JRT
UBE4B_HUMAN_3L1X
UBR5_HUMAN_1I2T
VILI_CHICK_1YU5
Reconstructed ΔG of a double mutant
ΔG of a double mutant
epistasis
no epistasis
Sequence count
1838 98
975 95
338 56
1811 257
625 69
271 64
401 30
543 55
360 1
1725 645
350 9
1199 106

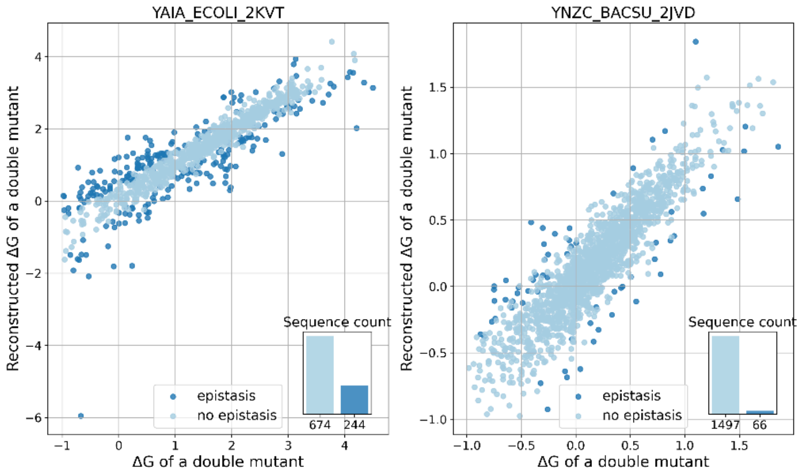


**Supplementary Figure 3.** Epistatic and non-epistatic genotypes for the Tsuboyama stability datasets for $N$ = 3: for each multi-mutant genotype, the x-axis shows its observed brightness and the y-axis shows the sum of the corresponding single-mutant brightness values. Both values are shown as raw brightness values (not log-transformed) and relative to the wildtype. Dark blue dots correspond to the epistatic genotypes, i.e. where the functional effect of a combination of mutations significantly (larger than the experimental error) differs from the linear combination of individual effects.

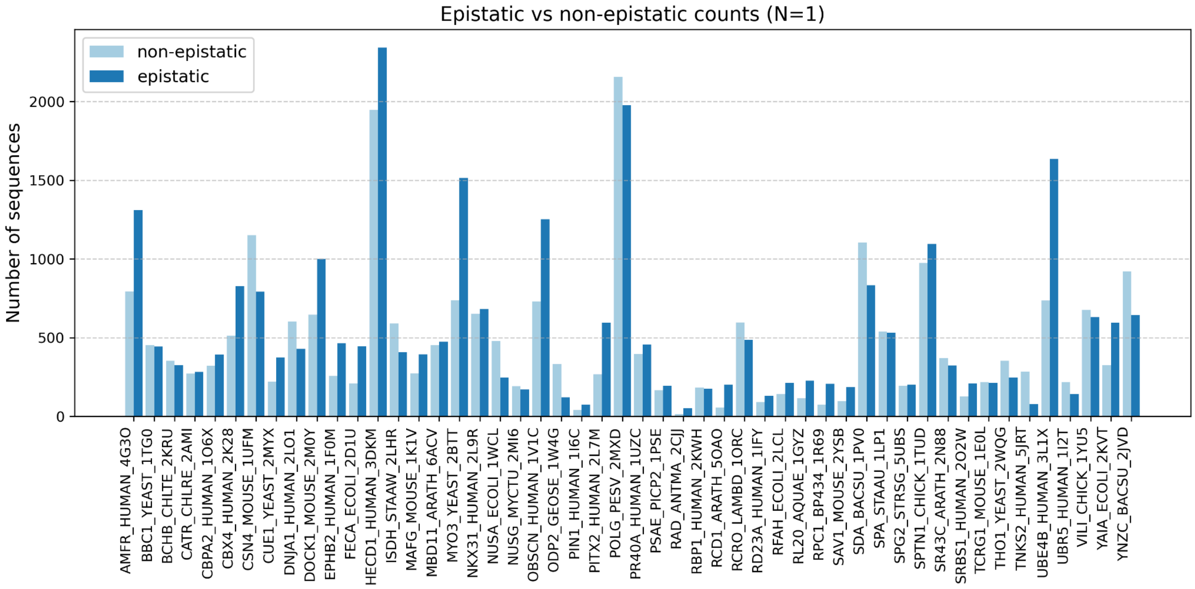
Epistatic vs non-epistatic counts (N=1)
non-epistatic
epistatic
Number of sequences
0
500
1000
1500
2000
AMFR_HUMAN_4G3O
BBC1_YEAST_1TG0
BCHB_CHLTE_2KRU
CATR_CHLRE_2AMI
CBPA2_HUMAN_1O6X
CBX4_HUMAN_2K28
CSN4_MOUSE_1UFM
CUE1_YEAST_2MYX
DNJA1_HUMAN_2LO1
DOCK1_MOUSE_2M0Y
EPHB2_HUMAN_1F0M
FECA_ECOLI_2D1U
HECD1_HUMAN_3DKM
ISDH_STAAW_2LHR
MAFG_MOUSE_1K1V
MBD11_ARATH_6ACV
MYO3_YEAST_2BTT
NKX31_HUMAN_2L9R
NUSA_ECOLI_1WCL
NUSG_MYCTU_2MI6
OBSCN_HUMAN_1V1C
ODP2_GEOSE_1W4G
PIN1_HUMAN_1I6C
PITX2_HUMAN_2L7M
POLG_PESV_2MXD
PR40A_HUMAN_1UZC
PSAE_PICP2_1PSE
RAD_ANTMA_2CJJ
RBP1_HUMAN_2KWH
RCD1_ARATH_5OAO
RCRO_LAMBD_1ORC
RD23A_HUMAN_1IFY
RFAH_ECOLI_2LCL
RL20_AQUAE_1GYZ
RPC1_BP434_1R69
SAV1_MOUSE_2YSB
SDA_BACSU_1PV0
SPA_STAAU_1LP1
SPG2_STRSG_5UBS
SPTN1_CHICK_1TUD
SR43C_ARATH_2N88
SRBS1_HUMAN_2O2W
TCRG1_MOUSE_1E0L
THO1_YEAST_2WQG
TNKS2_HUMAN_5JRT
UBE4B_HUMAN_3L1X
UBR5_HUMAN_1I2T
VILI_CHICK_1YU5
YAIA_ECOLI_2KVT
YNZC_BACSU_2JVD

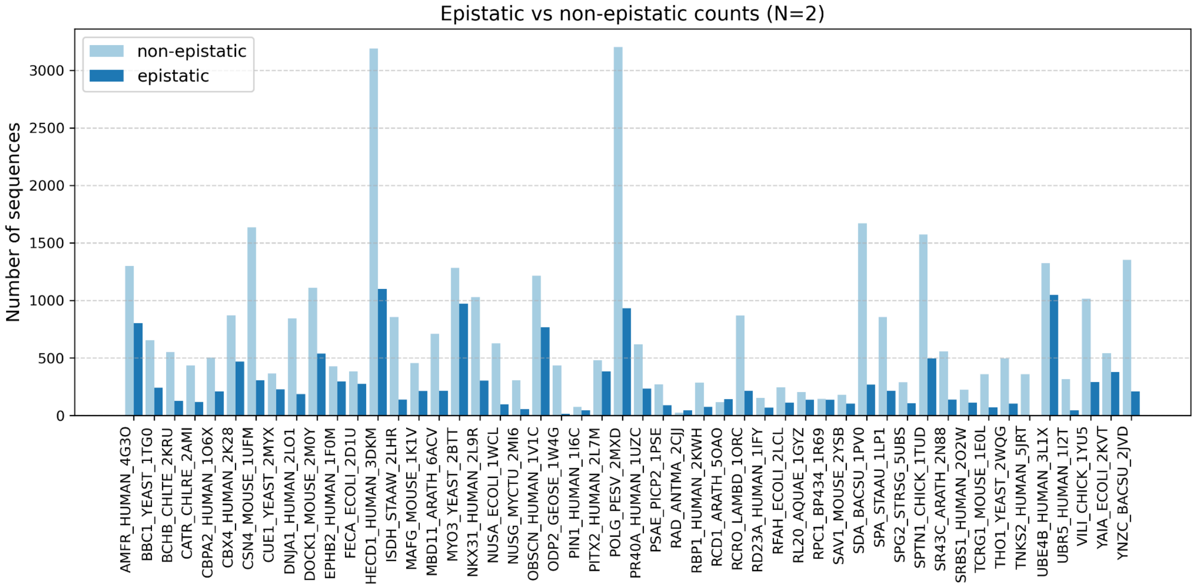
Epistatic vs non-epistatic counts (N=2)
non-epistatic
epistatic
Number of sequences
0
500
1000
1500
2000
2500
3000
AMFR_HUMAN_4G3O
BBC1_YEAST_1TG0
BCHB_CHLTE_2KRU
CATR_CHLRE_2AMI
CBPA2_HUMAN_1O6X
CBX4_HUMAN_2K28
CSN4_MOUSE_1UFM
CUE1_YEAST_2MYX
DNJA1_HUMAN_2LO1
DOCK1_MOUSE_2M0Y
EPHB2_HUMAN_1F0M
FECA_ECOLI_2D1U
HECD1_HUMAN_3DKM
ISDH_STAAW_2LHR
MAFG_MOUSE_1K1V
MBD11_ARATH_6ACV
MYO3_YEAST_2BTT
NKX31_HUMAN_2L9R
NUSA_ECOLI_1WCL
NUSG_MYCTU_2MI6
OBSCN_HUMAN_1V1C
ODP2_GEOSE_1W4G
PIN1_HUMAN_1I6C
PITX2_HUMAN_2L7M
POLG_PESV_2MXD
PR40A_HUMAN_1UZC
PSAE_PICP2_1PSE
RAD_ANTMA_2CJJ
RBP1_HUMAN_2KWH
RCD1_ARATH_5OAO
RCRO_LAMBD_1ORC
RD23A_HUMAN_1IFY
RFAH_ECOLI_2LCL
RL20_AQUAE_1GYZ
RPC1_BP434_1R69
SAV1_MOUSE_2YSB
SDA_BACSU_1PV0
SPA_STAAU_1LP1
SPG2_STRSG_5UBS
SPTN1_CHICK_1TUD
SR43C_ARATH_2N88
SRBS1_HUMAN_2O2W
TCRG1_MOUSE_1E0L
THO1_YEAST_2WQG
TNKS2_HUMAN_5JRT
UBE4B_HUMAN_3L1X
UBR5_HUMAN_1I2T
VILI_CHICK_1YU5
YAIA_ECOLI_2KVT
YNZC_BACSU_2JVD

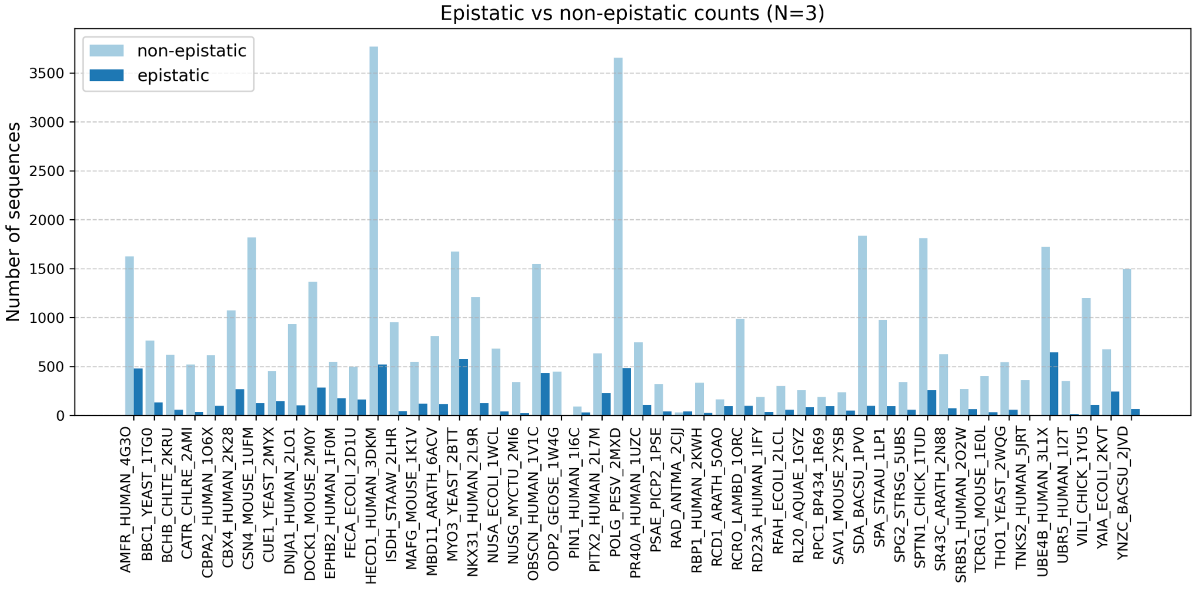
Epistatic vs non-epistatic counts (N=3)
non-epistatic
epistatic
Number of sequences
0
500
1000
1500
2000
2500
3000
3500
AMFR_HUMAN_4G3O
BBC1_YEAST_1TG0
BCHB_CHLTE_2KRU
CATR_CHLRE_2AMI
CBPA2_HUMAN_1O6X
CBX4_HUMAN_2K28
CSN4_MOUSE_1UFM
CUE1_YEAST_2MYX
DNJA1_HUMAN_2LO1
DOCK1_MOUSE_2M0Y
EPHB2_HUMAN_1F0M
FECA_ECOLI_2D1U
HECD1_HUMAN_3DKM
ISDH_STAAW_2LHR
MAFG_MOUSE_1K1V
MBD11_ARATH_6ACV
MYO3_YEAST_2BTT
NKX31_HUMAN_2L9R
NUSA_ECOLI_1WCL
NUSG_MYCTU_2MI6
OBSCN_HUMAN_1V1C
ODP2_GEOSE_1W4G
PIN1_HUMAN_1I6C
PITX2_HUMAN_2L7M
POLG_PESV_2MXD
PR40A_HUMAN_1UZC
PSAE_PICP2_1PSE
RAD_ANTMA_2CJJ
RBP1_HUMAN_2KWH
RCD1_ARATH_5OAO
RCRO_LAMBD_1ORC
RD23A_HUMAN_1IFY
RFAH_ECOLI_2LCL
RL20_AQUAE_1GYZ
RPC1_BP434_1R69
SAV1_MOUSE_2YSB
SDA_BACSU_1PV0
SPA_STAAU_1LP1
SPG2_STRSG_5UBS
SPTN1_CHICK_1TUD
SR43C_ARATH_2N88
SRBS1_HUMAN_2O2W
TCRG1_MOUSE_1E0L
THO1_YEAST_2WQG
TNKS2_HUMAN_5JRT
UBE4B_HUMAN_3L1X
UBR5_HUMAN_1I2T
VILI_CHICK_1YU5
YAIA_ECOLI_2KVT
YNZC_BACSU_2JVD

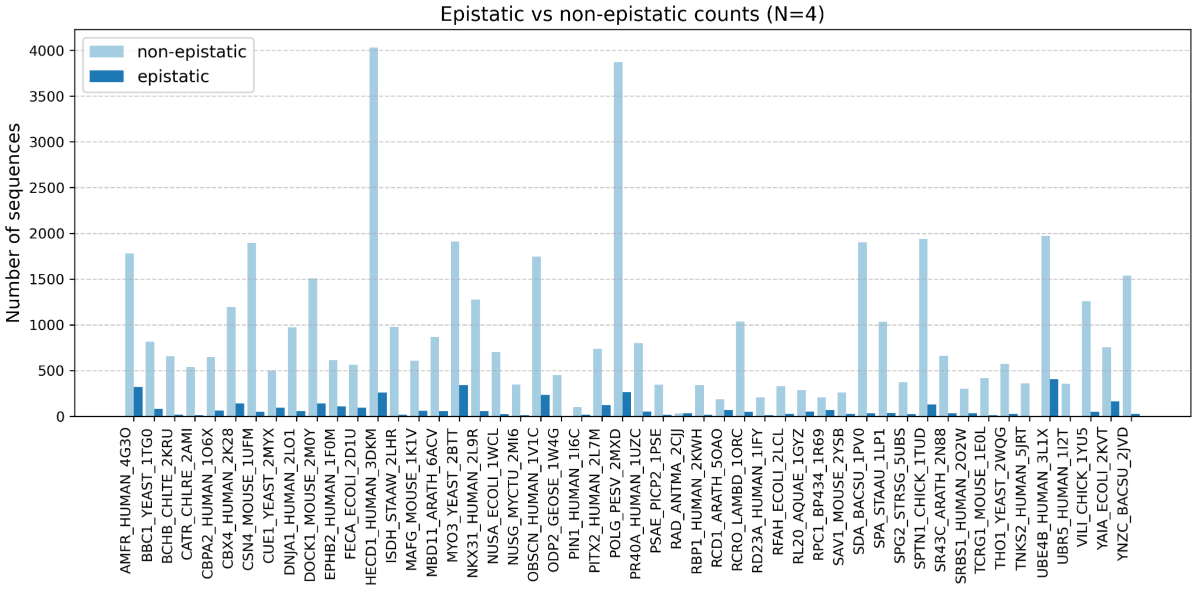
Epistatic vs non-epistatic counts (N=4)
non-epistatic
epistatic
Number of sequences
0
500
1000
1500
2000
2500
3000
3500
4000
AMFR_HUMAN_4G3O
BBC1_YEAST_1TG0
BCHB_CHLTE_2KRU
CATR_CHLRE_2AMI
CBPA2_HUMAN_1O6X
CBX4_HUMAN_2K28
CSN4_MOUSE_1UFM
CUE1_YEAST_2MYX
DNJA1_HUMAN_2LO1
DOCK1_MOUSE_2M0Y
EPHB2_HUMAN_1F0M
FECA_ECOLI_2D1U
HECD1_HUMAN_3DKM
ISDH_STAAW_2LHR
MAFG_MOUSE_1K1V
MBD11_ARATH_6ACV
MYO3_YEAST_2BTT
NKX31_HUMAN_2L9R
NUSA_ECOLI_1WCL
NUSG_MYCTU_2MI6
OBSCN_HUMAN_1V1C
ODP2_GEOSE_1W4G
PIN1_HUMAN_1I6C
PITX2_HUMAN_2L7M
POLG_PESV_2MXD
PR40A_HUMAN_1UZC
PSAE_PICP2_1PSE
RAD_ANTMA_2CJJ
RBP1_HUMAN_2KWH
RCD1_ARATH_5OAO
RCRO_LAMBD_1ORC
RD23A_HUMAN_1IFY
RFAH_ECOLI_2LCL
RL20_AQUAE_1GYZ
RPC1_BP434_1R69
SAV1_MOUSE_2YSB
SDA_BACSU_1PV0
SPA_STAAU_1LP1
SPG2_STRSG_5UBS
SPTN1_CHICK_1TUD
SR43C_ARATH_2N88
SRBS1_HUMAN_2O2W
TCRG1_MOUSE_1E0L
THO1_YEAST_2WQG
TNKS2_HUMAN_5JRT
UBE4B_HUMAN_3L1X
UBR5_HUMAN_1I2T
VILI_CHICK_1YU5
YAIA_ECOLI_2KVT
YNZC_BACSU_2JVD

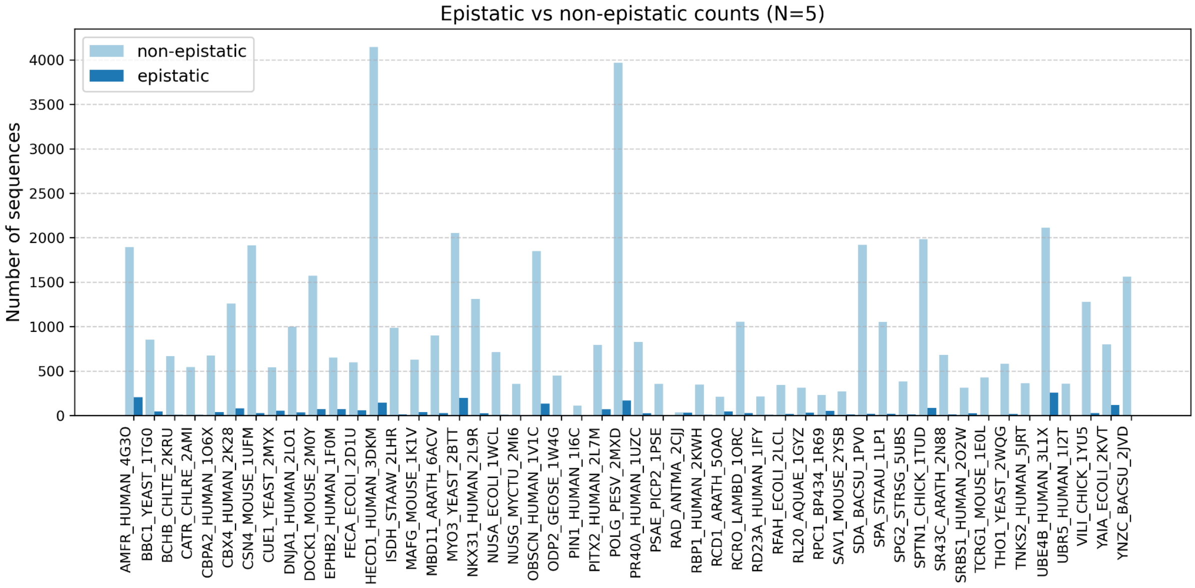
Epistatic vs non-epistatic counts (N=5)
non-epistatic
epistatic
Number of sequences
0
500
1000
1500
2000
2500
3000
3500
4000
AMFR_HUMAN_4G3O
BBC1_YEAST_1TG0
BCHB_CHLTE_2KRU
CATR_CHLRE_2AMI
CBPA2_HUMAN_1O6X
CBX4_HUMAN_2K28
CSN4_MOUSE_1UFM
CUE1_YEAST_2MYX
DNJA1_HUMAN_2LO1
DOCK1_MOUSE_2M0Y
EPHB2_HUMAN_1F0M
FECA_ECOLI_2D1U
HECD1_HUMAN_3DKM
ISDH_STAAW_2LHR
MAFG_MOUSE_1K1V
MBD11_ARATH_6ACV
MYO3_YEAST_2BTT
NKX31_HUMAN_2L9R
NUSA_ECOLI_1WCL
NUSG_MYCTU_2MI6
OBSCN_HUMAN_1V1C
ODP2_GEOSE_1W4G
PIN1_HUMAN_1I6C
PITX2_HUMAN_2L7M
POLG_PESV_2MXD
PR40A_HUMAN_1UZC
PSAE_PICP2_1PSE
RAD_ANTMA_2CJJ
RBP1_HUMAN_2KWH
RCD1_ARATH_5OAO
RCRO_LAMBD_1ORC
RD23A_HUMAN_1IFY
RFAH_ECOLI_2LCL
RL20_AQUAE_1GYZ
RPC1_BP434_1R69
SAV1_MOUSE_2YSB
SDA_BACSU_1PV0
SPA_STAAU_1LP1
SPG2_STRSG_5UBS
SPTN1_CHICK_1TUD
SR43C_ARATH_2N88
SRBS1_HUMAN_2O2W
TCRG1_MOUSE_1E0L
THO1_YEAST_2WQG
TNKS2_HUMAN_5JRT
UBE4B_HUMAN_3L1X
UBR5_HUMAN_1I2T
VILI_CHICK_1YU5
YAIA_ECOLI_2KVT
YNZC_BACSU_2JVD

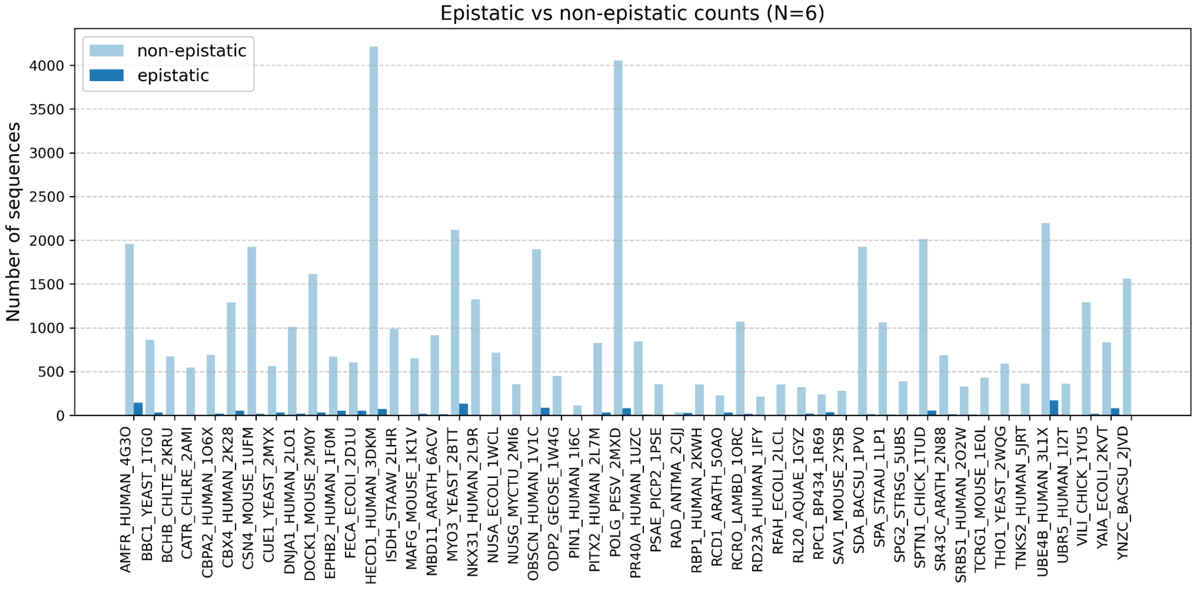
Epistatic vs non-epistatic counts (N=6)
non-epistatic
epistatic
Number of sequences
0
500
1000
1500
2000
2500
3000
3500
4000
AMFR_HUMAN_4G3O
BBC1_YEAST_1TG0
BCHB_CHLTE_2KRU
CATR_CHLRE_2AMI
CBPA2_HUMAN_1O6X
CBX4_HUMAN_2K28
CSN4_MOUSE_1UFM
CUE1_YEAST_2MYX
DNJA1_HUMAN_2LO1
DOCK1_MOUSE_2M0Y
EPHB2_HUMAN_1F0M
FECA_ECOLI_2D1U
HECD1_HUMAN_3DKM
ISDH_STAAW_2LHR
MAFG_MOUSE_1K1V
MBD11_ARATH_6ACV
MYO3_YEAST_2BTT
NKX31_HUMAN_2L9R
NUSA_ECOLI_1WCL
NUSG_MYCTU_2MI6
OBSCN_HUMAN_1V1C
ODP2_GEOSE_1W4G
PIN1_HUMAN_1I6C
PITX2_HUMAN_2L7M
POLG_PESV_2MXD
PR40A_HUMAN_1UZC
PSAE_PICP2_1PSE
RAD_ANTMA_2CJJ
RBP1_HUMAN_2KWH
RCD1_ARATH_5OAO
RCRO_LAMBD_1ORC
RD23A_HUMAN_1IFY
RFAH_ECOLI_2LCL
RL20_AQUAE_1GYZ
RPC1_BP434_1R69
SAV1_MOUSE_2YSB
SDA_BACSU_1PV0
SPA_STAAU_1LP1
SPG2_STRSG_5UBS
SPTN1_CHICK_1TUD
SR43C_ARATH_2N88
SRBS1_HUMAN_2O2W
TCRG1_MOUSE_1E0L
THO1_YEAST_2WQG
TNKS2_HUMAN_5JRT
UBE4B_HUMAN_3L1X
UBR5_HUMAN_1I2T
VILI_CHICK_1YU5
YAIA_ECOLI_2KVT
YNZC_BACSU_2JVD

**Supplementary Figure 4.** Barplots of the number of epistatic and non-epistatic sequences in the Tsuboyama datasets, depending on the selection of *N*.

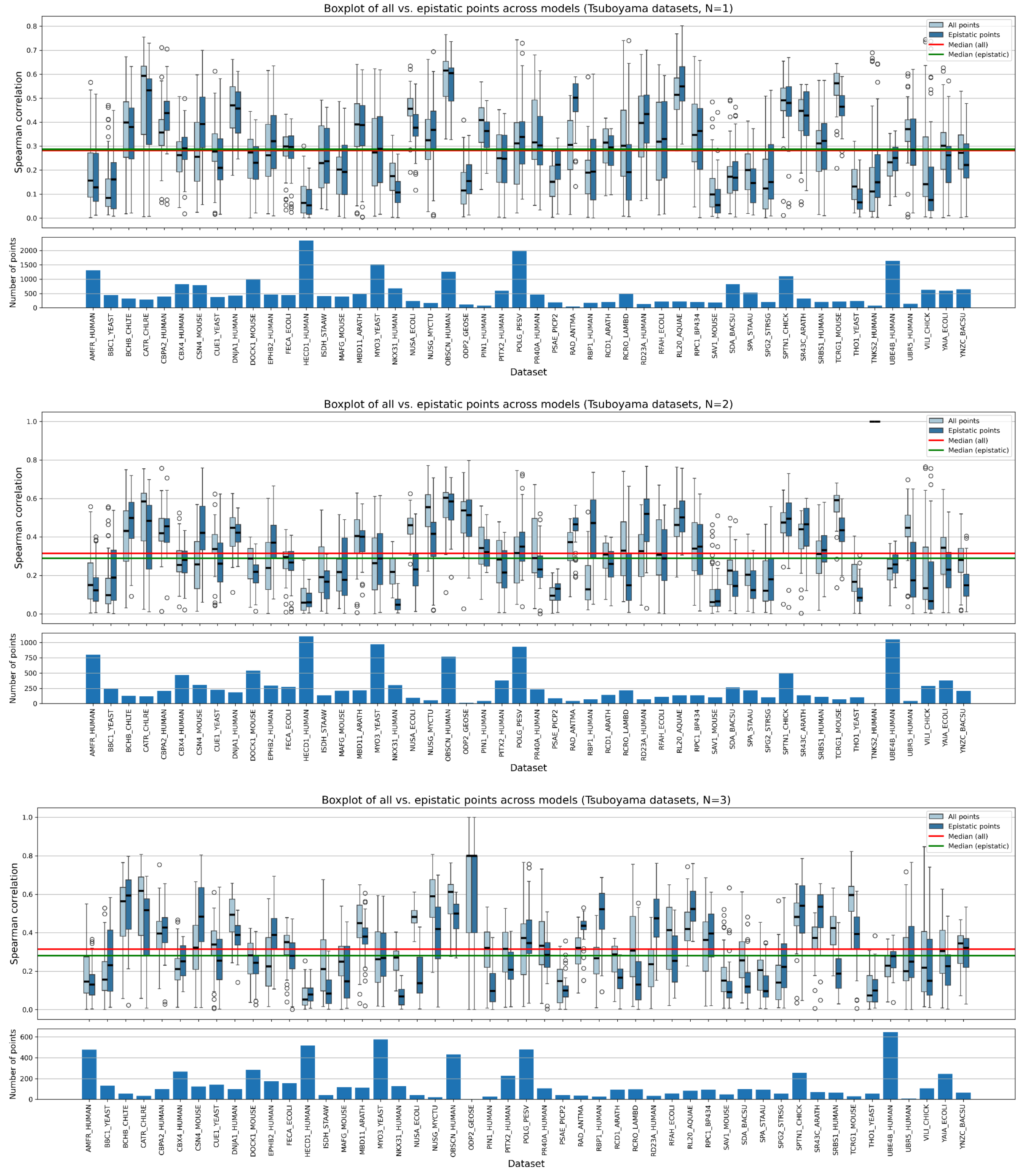

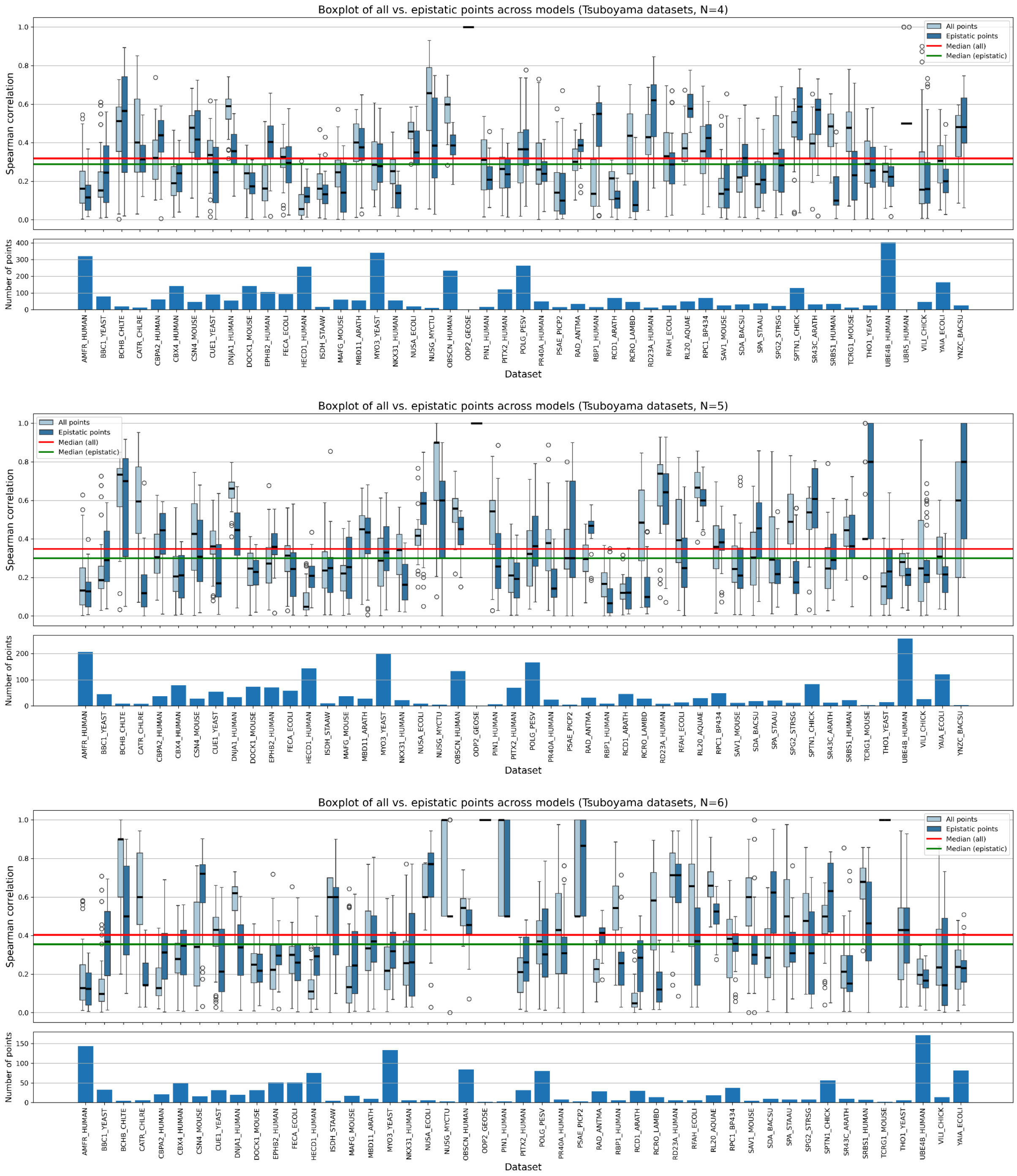


**Supplementary Figure 5.** Performance of zero-shot models for the stability datasets, depending on the selection of *N*. Correlations are computed separately for epistatic genotypes (dark blue) and a size-matched random sample of all multi-mutant genotypes (light blue). The lower panel shows the number of epistatic genotypes for each dataset.

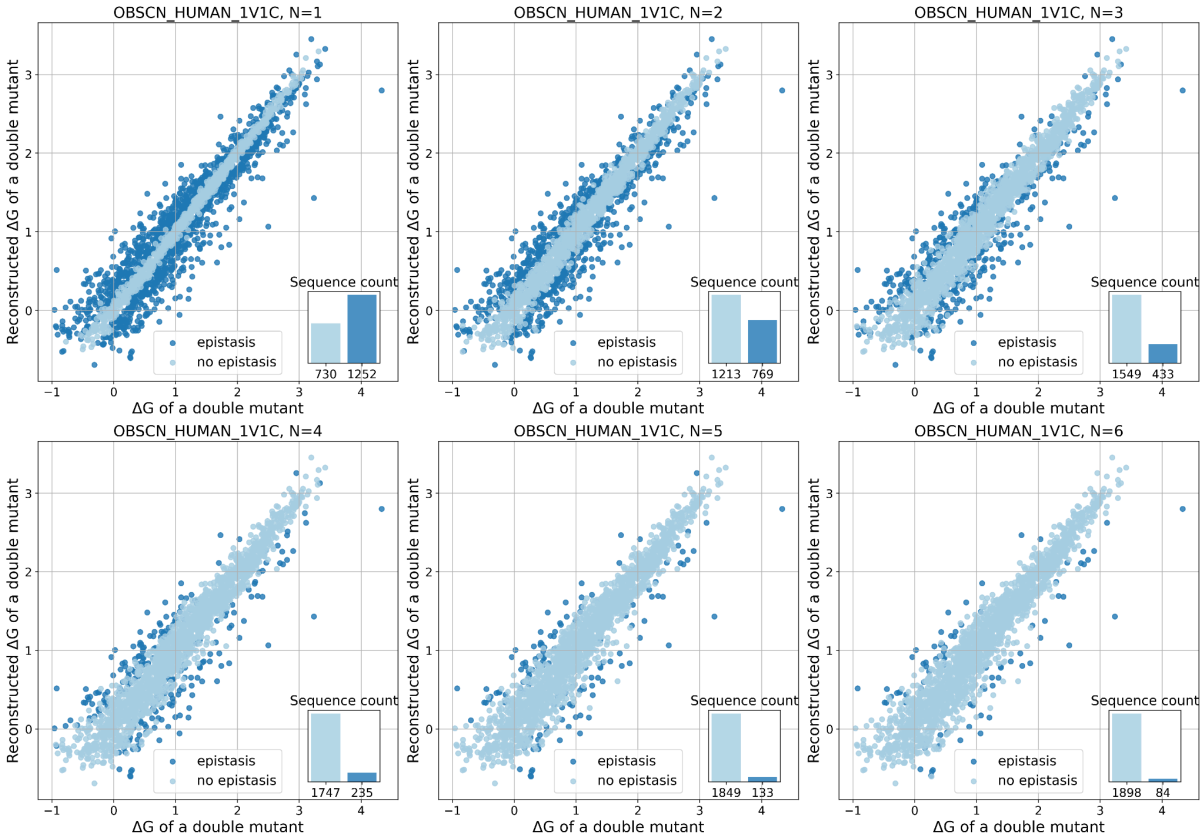


**Supplementary Figure 6.** Epistatic and non-epistatic genotypes on the example of OBSCN_HUMAN_1V1C from the Tsuboyama dataset, depending on the selection of N. Results for N=1, 2, 3 are demonstrated, as well as the sequence count: epistatic and non-epistatic. For each multi-mutant genotype, the x-axis shows its observed brightness and the y-axis shows the sum of the corresponding single-mutant brightness values. Both values are shown as raw brightness values (not log-transformed) and relative to the wildtype. Dark blue dots correspond to the epistatic genotypes, i.e. where the functional effect of a combination of mutations significantly (larger than the experimental error) differs from the linear combination of individual effects.

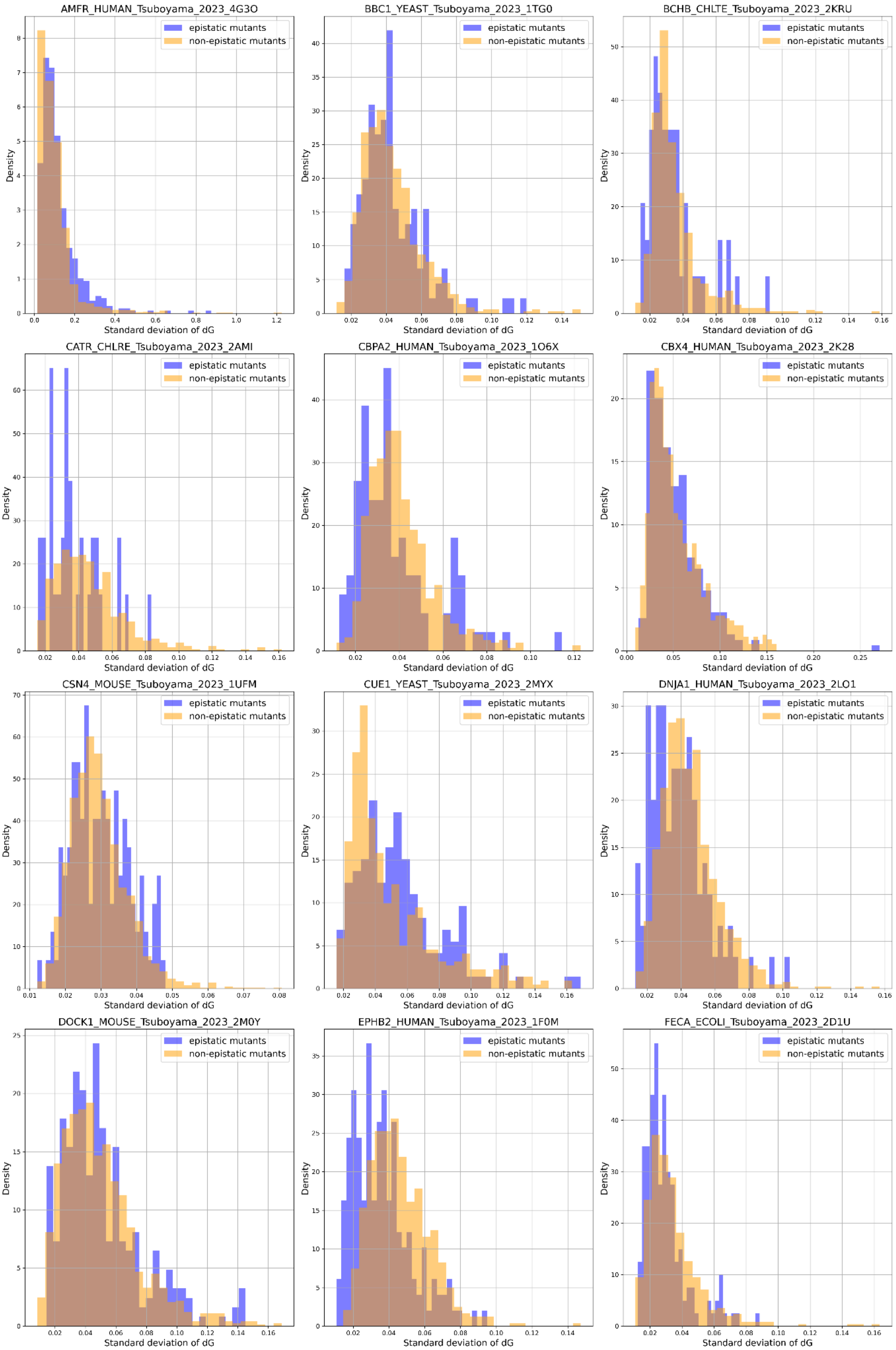
AMFR_HUMAN_Tsuboyama_2023_4G3O
BBC1_YEAST_Tsuboyama_2023_1TG0
BCHB_CHLTE_Tsuboyama_2023_2KRU
CATR_CHLRE_Tsuboyama_2023_2AMI
CBPA2_HUMAN_Tsuboyama_2023_1O6X
CBX4_HUMAN_Tsuboyama_2023_2K28
CSN4_MOUSE_Tsuboyama_2023_1UFM
CUE1_YEAST_Tsuboyama_2023_2MYX
DNJA1_HUMAN_Tsuboyama_2023_2LO1
DOCK1_MOUSE_Tsuboyama_2023_2M0Y
EPHB2_HUMAN_Tsuboyama_2023_1F0M
FECA_ECOLI_Tsuboyama_2023_2D1U
epistatic mutants
non-epistatic mutants
Density
Standard deviation of dG

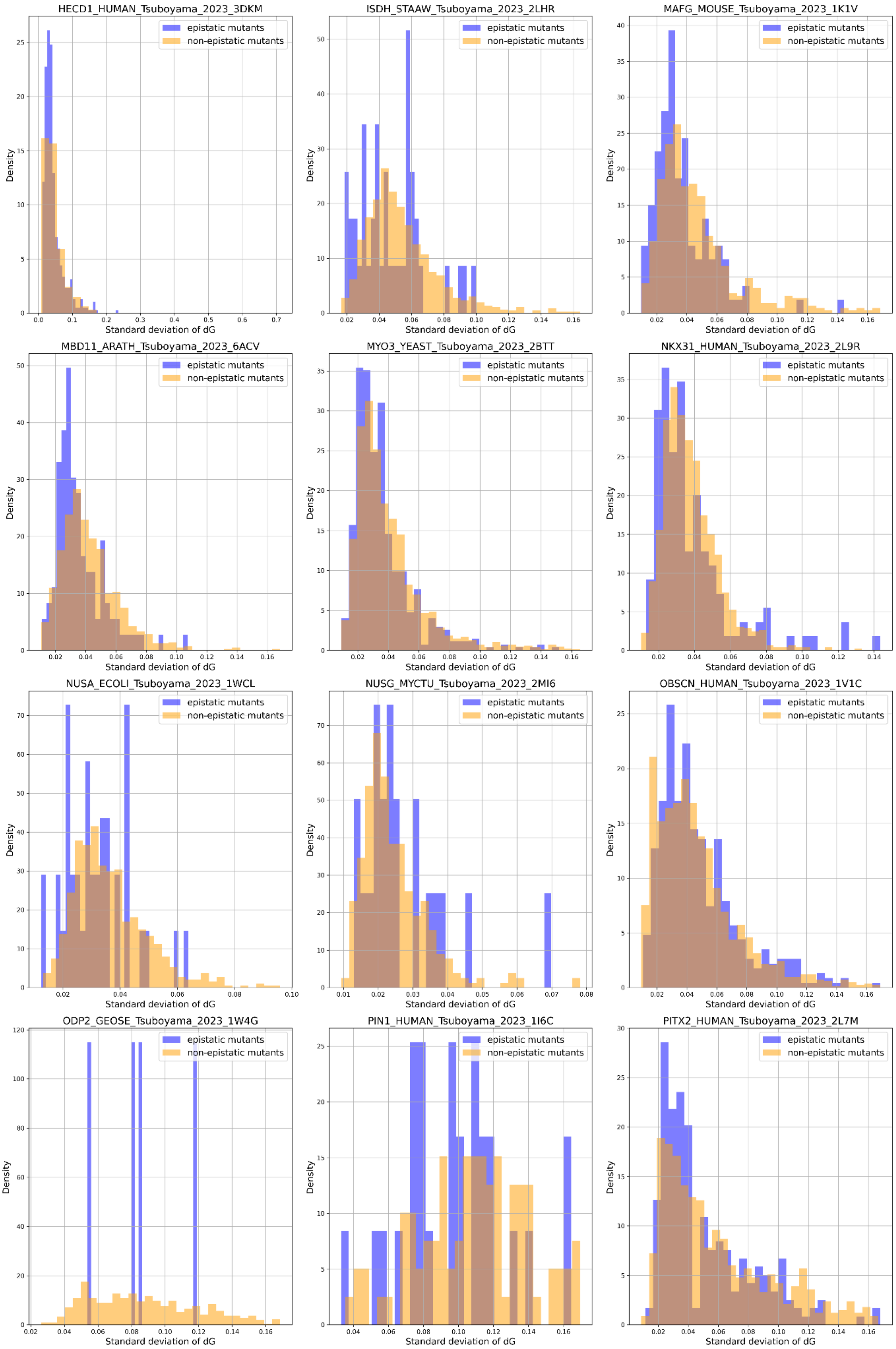
HECD1_HUMAN_Tsuboyama_2023_3DKM
ISDH_STAAW_Tsuboyama_2023_2LHR
MAFG_MOUSE_Tsuboyama_2023_1K1V
MBD11_ARATH_Tsuboyama_2023_6ACV
MYO3_YEAST_Tsuboyama_2023_2BTT
NKX31_HUMAN_Tsuboyama_2023_2L9R
NUSA_ECOLI_Tsuboyama_2023_1WCL
NUSG_MYCTU_Tsuboyama_2023_2MI6
OBSCN_HUMAN_Tsuboyama_2023_1V1C
ODP2_GEOSE_Tsuboyama_2023_1W4G
PIN1_HUMAN_Tsuboyama_2023_1I6C
PITX2_HUMAN_Tsuboyama_2023_2L7M
epistatic mutants
non-epistatic mutants
Density
Standard deviation of dG

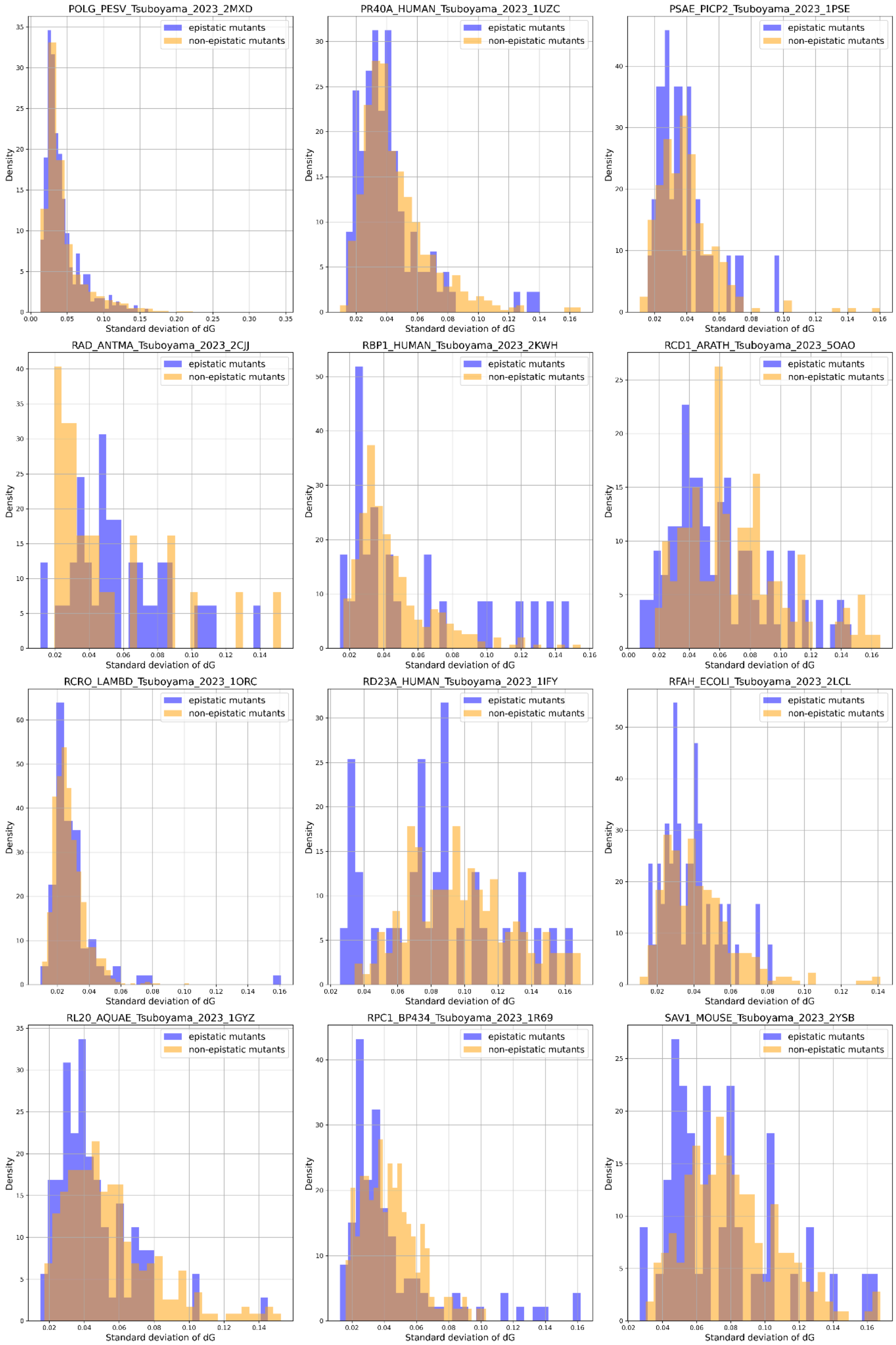
POLG_PESV_Tsuboyama_2023_2MXD
PR40A_HUMAN_Tsuboyama_2023_1UZC
PSAE_PICP2_Tsuboyama_2023_1PSE
RAD_ANTMA_Tsuboyama_2023_2CJJ
RBP1_HUMAN_Tsuboyama_2023_2KWH
RCD1_ARATH_Tsuboyama_2023_5OAO
RCRO_LAMBD_Tsuboyama_2023_1ORC
RD23A_HUMAN_Tsuboyama_2023_1IFY
RFAH_ECOLI_Tsuboyama_2023_2LCL
RL20_AQUAE_Tsuboyama_2023_1GYZ
RPC1_BP434_Tsuboyama_2023_1R69
SAV1_MOUSE_Tsuboyama_2023_2YSB
epistatic mutants
non-epistatic mutants
Density
Standard deviation of dG

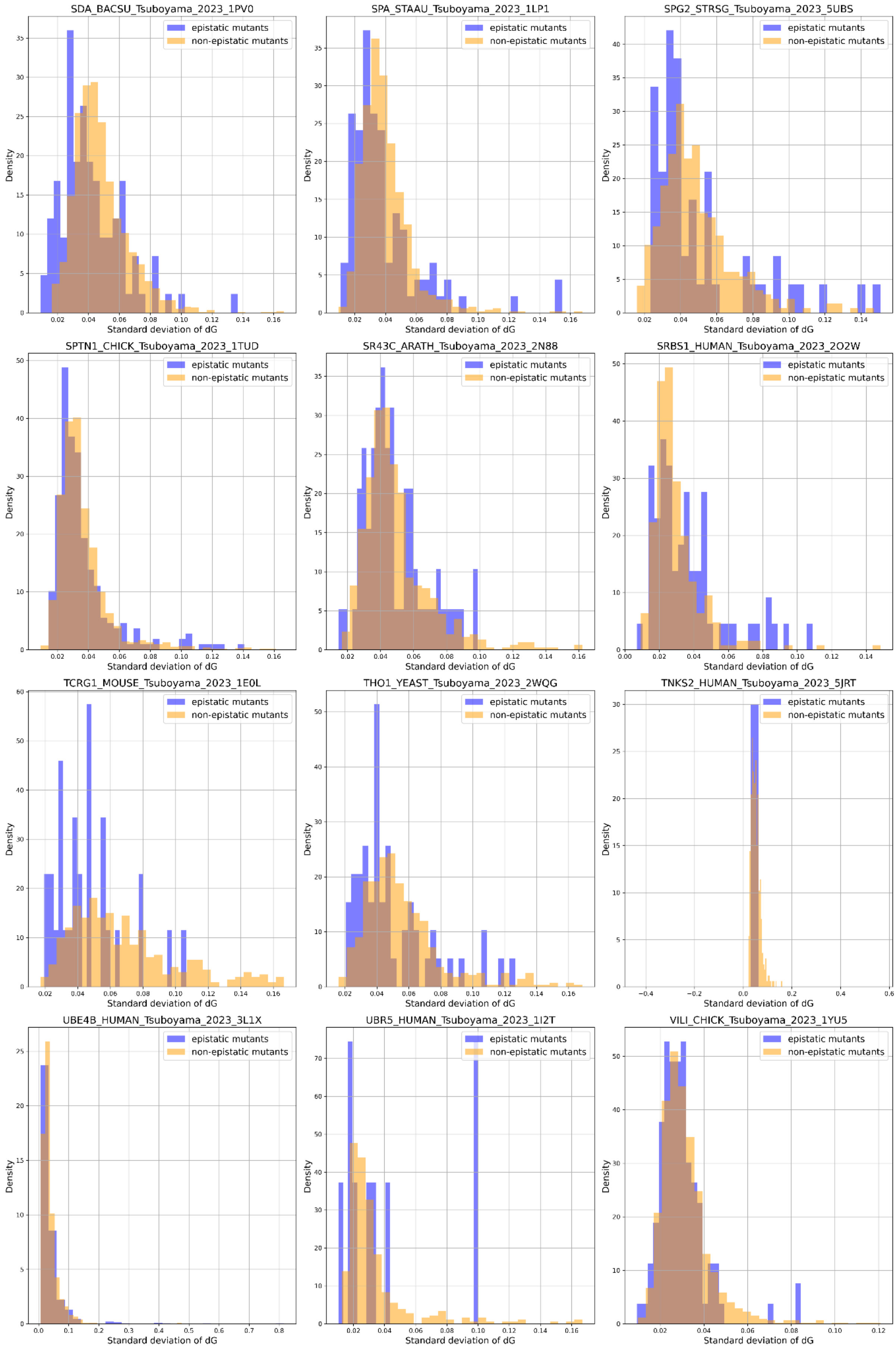

SDA_BACSU_Tsuboyama_2023_1PV0
SPA_STAAU_Tsuboyama_2023_1LP1
SPG2_STRSG_Tsuboyama_2023_5UBS
SPTN1_CHICK_Tsuboyama_2023_1TUD
SR43C_ARATH_Tsuboyama_2023_2N88
SRBS1_HUMAN_Tsuboyama_2023_2O2W
TCRG1_MOUSE_Tsuboyama_2023_1E0L
THO1_YEAST_Tsuboyama_2023_2WQG
TNKS2_HUMAN_Tsuboyama_2023_5JRT
UBE4B_HUMAN_Tsuboyama_2023_3L1X
UBR5_HUMAN_Tsuboyama_2023_1I2T
VILI_CHICK_Tsuboyama_2023_1YU5
epistatic mutants
non-epistatic mutants
Density
Standard deviation of dG

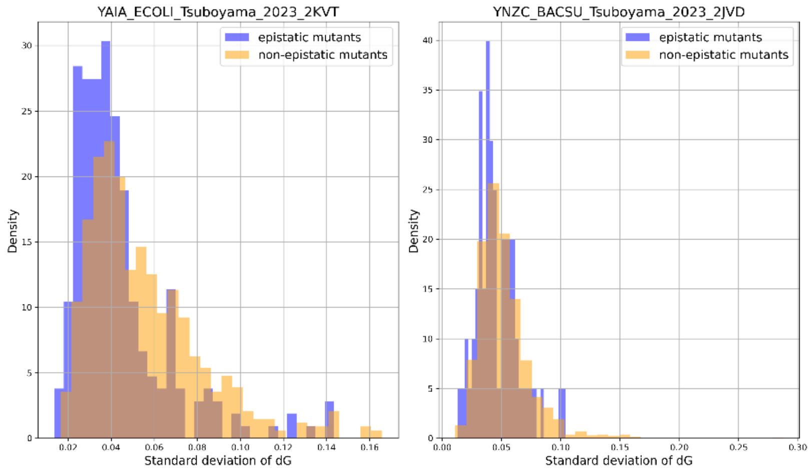


**Supplementary Figure 7.** The distribution of experimental errors in epistatic (dark blue) and non-epistatic (orange) genotypes for the Tsuboyama datasets. Derivation of the standard deviation of dG is explained in Methods. Observed epistasis cannot be attributed to larger experimental errors, and hence is a genuine biological effect.

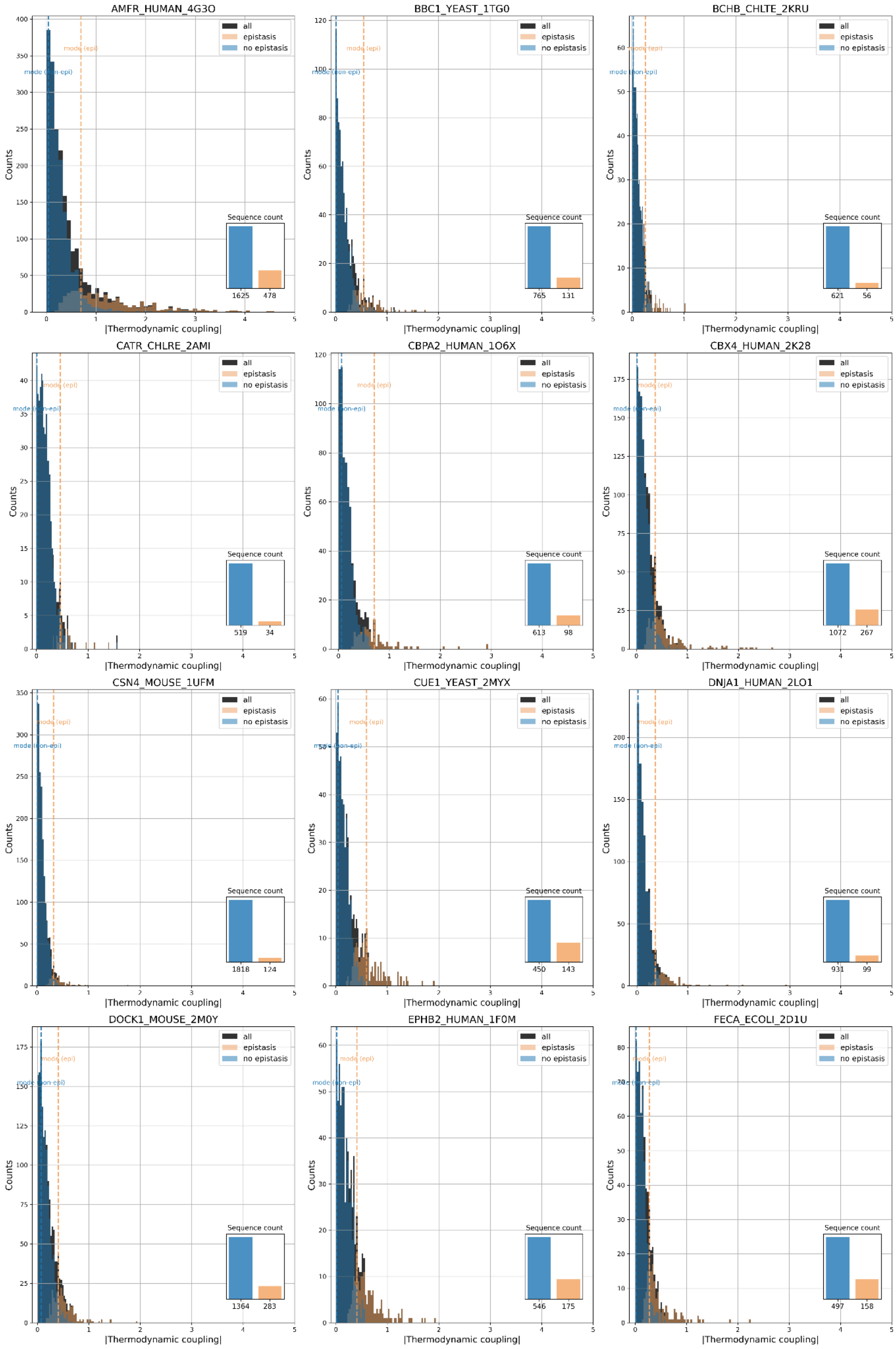
AMFR_HUMAN_4G3O
all
epistasis
no epistasis
mode (epi)
mode (non-epi)
Counts
|Thermodynamic coupling|
Sequence count
1625
478
BBC1_YEAST_1TG0
Sequence count
765
131
BCHB_CHLTE_2KRU
Sequence count
621
56
CATR_CHLRE_2AMI
Sequence count
519
34
CBPA2_HUMAN_1O6X
Sequence count
613
98
CBX4_HUMAN_2K28
Sequence count
1072
267
CSN4_MOUSE_1UFM
Sequence count
1818
124
CUE1_YEAST_2MYX
Sequence count
450
143
DNJA1_HUMAN_2LO1
Sequence count
931
99
DOCK1_MOUSE_2M0Y
Sequence count
1364
283
EPHB2_HUMAN_1F0M
Sequence count
546
175
FECA_ECOLI_2D1U
Sequence count
497
158

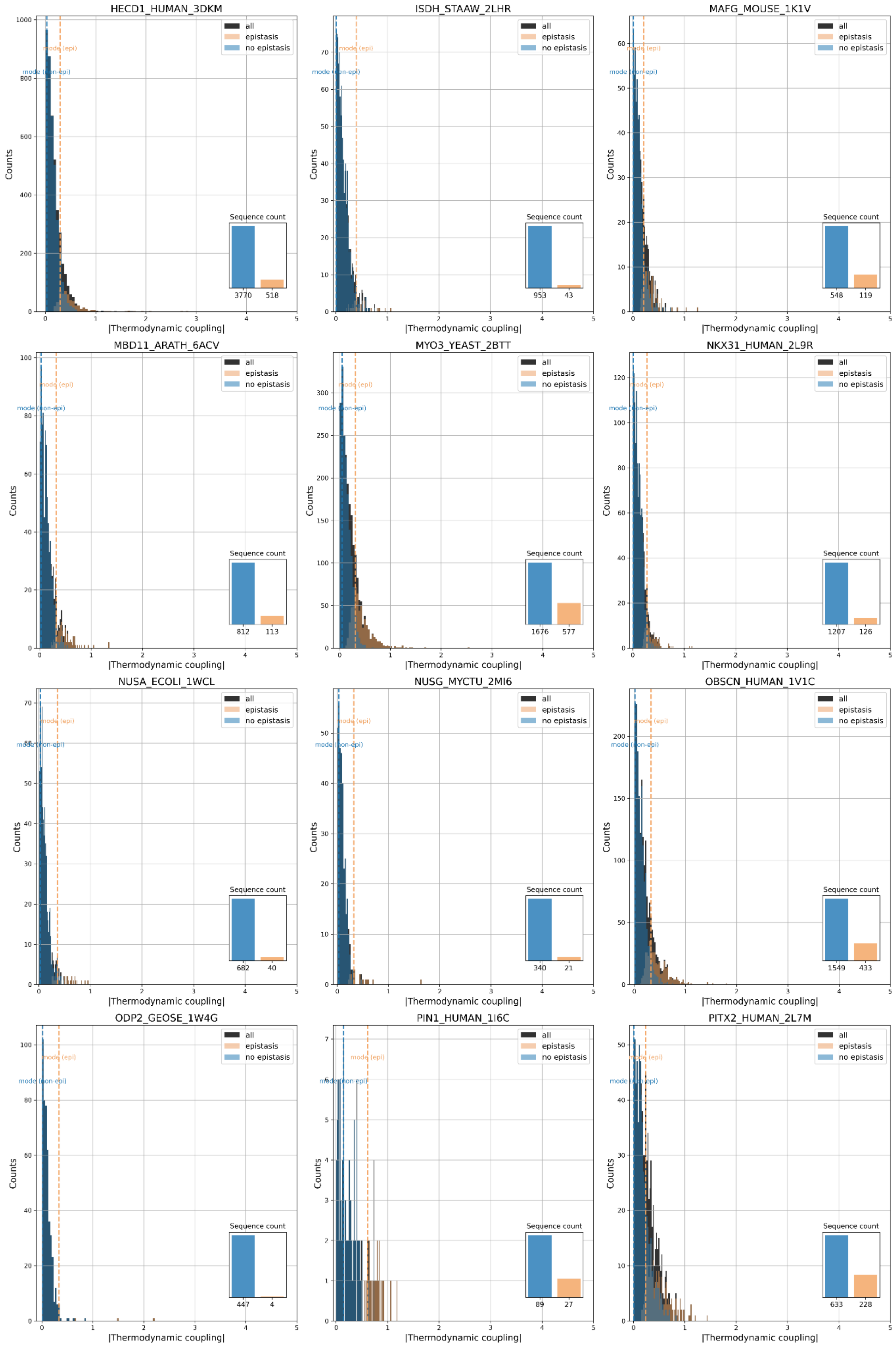
HECD1_HUMAN_3DKM
all
epistasis
no epistasis
mode (epi)
mode (non-epi)
Counts
Sequence count
3770
518
|Thermodynamic coupling|
ISDH_STAAW_2LHR
953
43
MAFG_MOUSE_1K1V
548
119
MBD11_ARATH_6ACV
812
113
MYO3_YEAST_2BTT
1676
577
NKX31_HUMAN_2L9R
1207
126
NUSA_ECOLI_1WCL
682
40
NUSG_MYCTU_2MI6
340
21
OBSCN_HUMAN_1V1C
1549
433
ODP2_GEOSE_1W4G
447
4
PIN1_HUMAN_1I6C
89
27
PITX2_HUMAN_2L7M
633
228

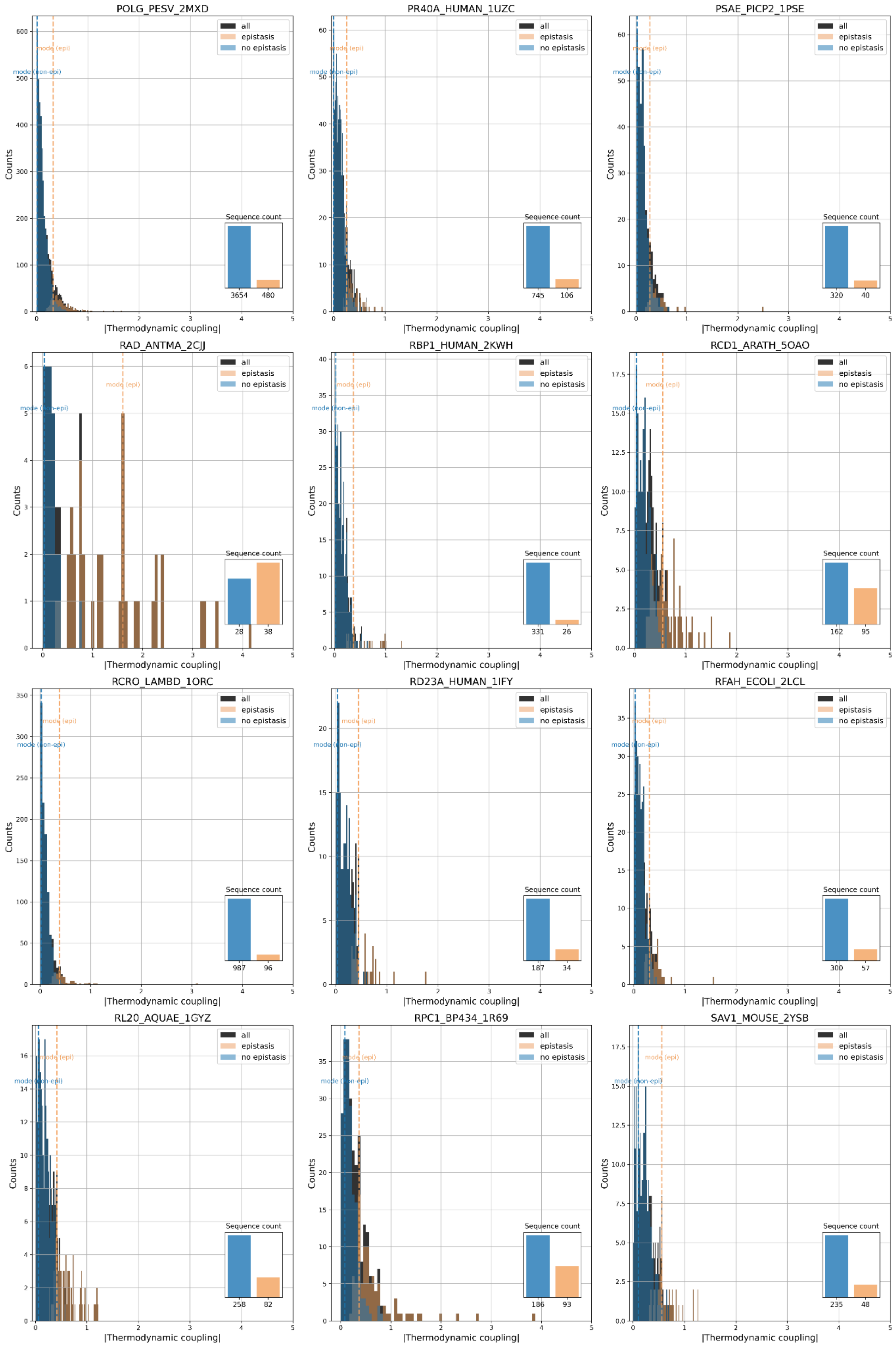

POLG_PESV_2MXD
all
epistasis
no epistasis
mode (epi)
mode (non-epi)
Counts
Sequence count
3654
480
|Thermodynamic coupling|
PR40A_HUMAN_1UZC
Sequence count
745
106
PSAE_PICP2_1PSE
Sequence count
320
40
RAD_ANTMA_2CJJ
Sequence count
28
38
RBP1_HUMAN_2KWH
Sequence count
331
26
RCD1_ARATH_5OAO
Sequence count
162
95
RCRO_LAMBD_1ORC
Sequence count
987
96
RD23A_HUMAN_1IFY
Sequence count
187
34
RFAH_ECOLI_2LCL
Sequence count
300
57
RL20_AQUAE_1GYZ
Sequence count
258
82
RPC1_BP434_1R69
Sequence count
186
93
SAV1_MOUSE_2YSB
Sequence count
235
48

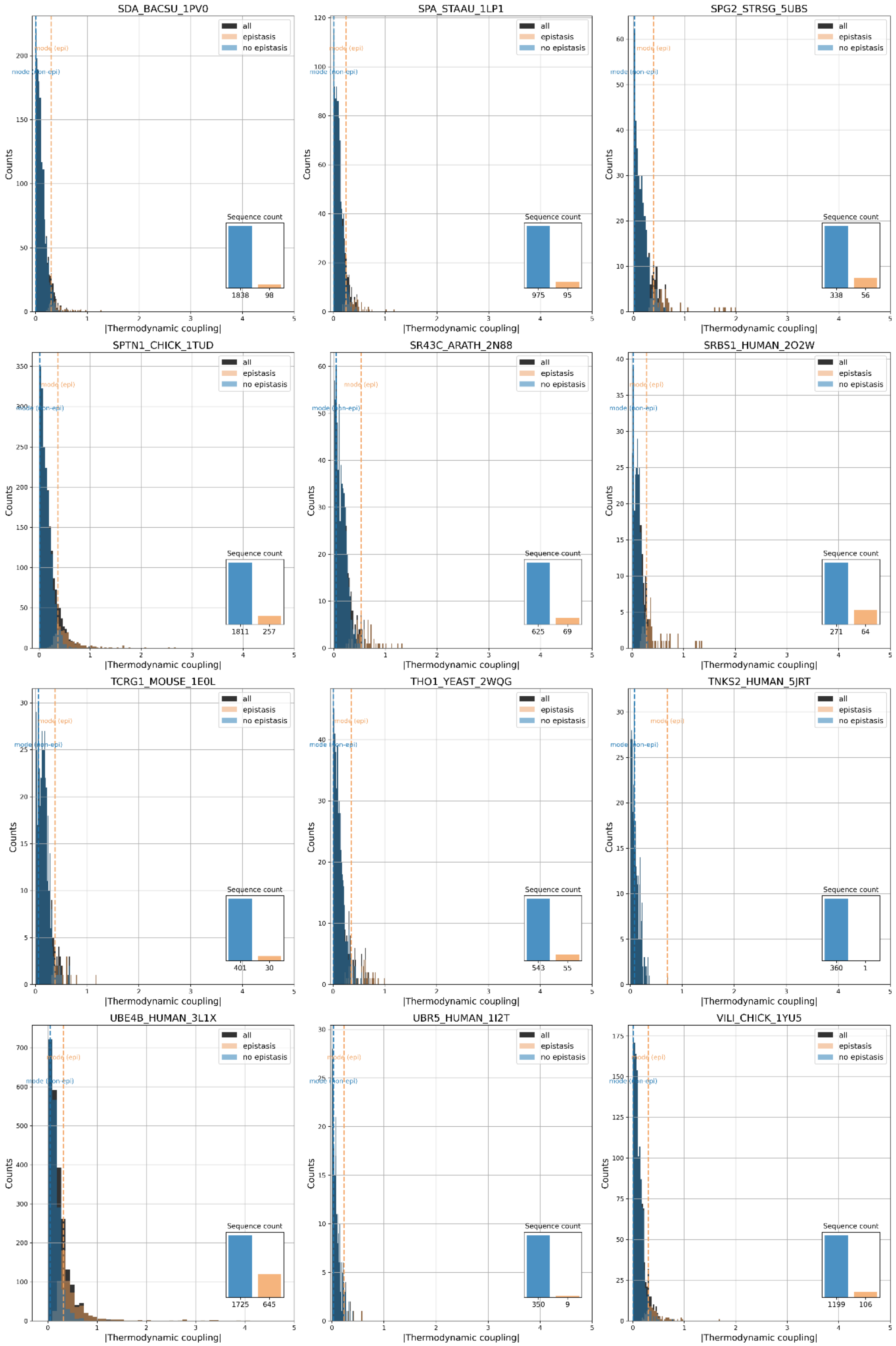
SDA_BACSU_1PV0
SPA_STAAU_1LP1
SPG2_STRSG_5UBS
SPTN1_CHICK_1TUD
SR43C_ARATH_2N88
SRBS1_HUMAN_2O2W
TCRG1_MOUSE_1E0L
THO1_YEAST_2WQG
TNKS2_HUMAN_5JRT
UBE4B_HUMAN_3L1X
UBR5_HUMAN_1I2T
VILI_CHICK_1YU5
all
epistasis
no epistasis
mode (epi)
mode (non-epi)
Counts
|Thermodynamic coupling|
Sequence count
1838
98
975
95
338
56
1811
257
625
69
271
64
401
30
543
55
360
1
1725
645
350
9
1199
106

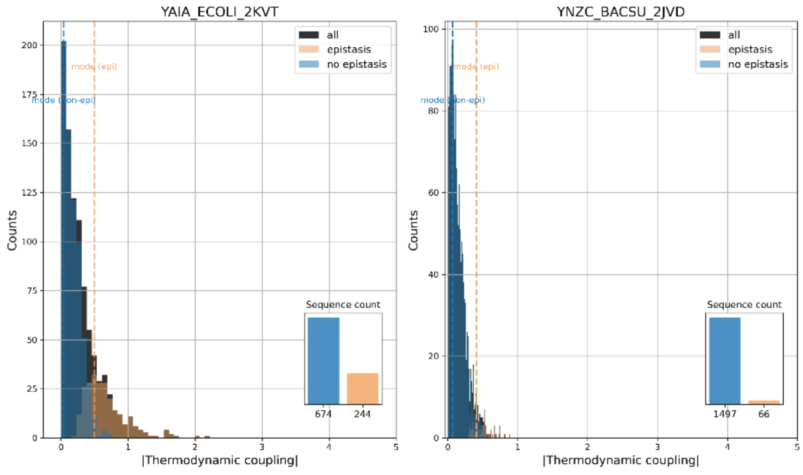


**Supplementary Figure 8.** Histograms of the absolute value of the thermodynamic couplings for epistatic and non-epistatic genotypes for the Tsuboyama datasets, as well as sequence counts. “all” stands for the combination of the two distributions.

**Supplementary Table 1.** For each Tsuboyama dataset, the number of epistatic sequences (“counts”) is shown.
https://github.com/kalininalab/epistasis_proteingym/blob/main/results/tables/intermediate/counts.csv
**Supplementary Table 2.** Best performing methods (see Methods for selection of best models), as well as their Spearman correlation for epistatic and “all” genotypes are shown for each Somermeyer dataset.
https://github.com/kalininalab/epistasis_proteingym/blob/main/results/tables/intermediate/models_evaluation/somermeyer_best_models.csv
**Supplementary Table 3.** Best performing methods (see Methods for selection of best models), as well as their Spearman correlation for epistatic and “all” genotypes are shown for each Tsuboyama dataset.
https://github.com/kalininalab/epistasis_proteingym/blob/main/results/tables/intermediate/models_evaluation/tsuboyama_best_models.csv

### Packages versions

```
channels:
  - conda-forge/label/cf201901
  - anaconda
  - https://repo.anaconda.com/pkgs/msys2
  - https://repo.anaconda.com/pkgs/free
  - defaults
dependencies:
  - _libgcc_mutex=0.1
  - _openmp_mutex=5.1
  - anyio=4.7.0
  - aom=3.6.0
  - argon2-cffi=21.3.0
```

- argon2-cffi-bindings=21.2.0
- asttokens=3.0.0
- async-lru=2.0.4
- attrs=24.3.0
- babel=2.16.0
- beautifulsoup4=4.13.4
- biopython=1.78
- blas=1.0
- bleach=6.2.0
- bottleneck=1.4.2
- brotli-python=1.0.9
- brotlicffi=1.0.9.2
- bzip2=1.0.8
- c-ares=1.19.1
- ca-certificates=2025.7.15
- cairo=1.16.0
- certifi=2025.8.3
- cffi=1.17.1
- charset-normalizer=3.3.2
- comm=0.2.1
- contourpy=1.3.1
- cudatoolkit=11.8.0
- cycler=0.11.0
- cyrus-sasl=2.1.28
- dav1d=1.2.1
- debugpy=1.8.11
- decorator=5.1.1
- defusedxml=0.7.1
- deprecated=1.2.13
- exceptiongroup=1.2.0
- executing=0.8.3
- expat=2.7.1
- filelock=3.17.0
- fontconfig=2.14.1
- fonttools=4.55.3
- freetype=2.13.3
- fribidi=1.0.10
- fsspec=2025.7.0
- gmp=6.3.0
- gmpy2=2.2.1
- graphite2=1.3.14
- h11=0.16.0
- harfbuzz=10.2.0
- httpcore=1.0.9
- httpx=0.28.1
- icu=73.1
- idna=3.7
- importlib-metadata=8.5.0

- ipykernel=6.29.5
- ipython=8.30.0
- ipywidgets=8.1.5
- jedi=0.19.2
- jinja2=3.1.6
- joblib=1.5.1
- jpeg=9e
- json5=0.9.25
- jsonschema=4.25.0
- jsonschema-specifications=2023.7.1
- jupyter=1.1.1
- jupyter-lsp=2.2.5
- jupyter_client=8.6.3
- jupyter_console=6.6.3
- jupyter_core=5.8.1
- jupyter_events=0.12.0
- jupyter_server=2.16.0
- jupyter_server_terminals=0.5.3
- jupyterlab=4.4.4
- jupyterlab_pygments=0.3.0
- jupyterlab_server=2.27.3
- jupyterlab_widgets=3.0.15
- kiwisolver=1.4.8
- lcms2=2.16
- ld_impl_linux-64=2.40
- lerc=4.0.0
- libabseil=20250127.0
- libavif=1.1.1
- libcups=2.4.2
- libcurl=8.14.1
- libdeflate=1.22
- libdrm=2.4.124
- libedit=3.1.20230828
- libegl=1.7.0
- libev=4.33
- libffi=3.4.4
- libgcc-ng=11.2.0
- libgfortran-ng=11.2.0
- libgfortran5=11.2.0
- libgl=1.7.0
- libglib=2.84.2
- libglvnd=1.7.0
- libglx=1.7.0
- libgomp=11.2.0
- libiconv=1.16
- libkrb5=1.21.3
- libllvm15=15.0.7
- libnghttp2=1.57.0

- libopenblas=0.3.29
- libpciaccess=0.18
- libpng=1.6.39
- libpq=17.4
- libprotobuf=5.29.3
- libsodium=1.0.18
- libssh2=1.11.1
- libstdcxx-ng=11.2.0
- libtiff=4.7.0
- libtorch=2.6.0
- libuuid=1.41.5
- libuv=1.48.0
- libwebp-base=1.3.2
- libxcb=1.17.0
- libxkbcommon=1.9.1
- libxml2=2.13.8
- lmdb=0.9.29
- lz4-c=1.9.4
- markupsafe=3.0.2
- matplotlib=3.10.0
- matplotlib-base=3.10.0
- matplotlib-inline=0.1.6
- mesalib=25.1.5
- mistune=3.1.2
- mpc=1.3.1
- mpfr=4.2.1
- mpmath=1.3.0
- mysql=8.4.0
- nbclient=0.10.2
- nbconvert=7.16.6
- nbconvert-core=7.16.6
- nbconvert-pandoc=7.16.6
- nbformat=5.10.4
- ncurses=6.5
- nest-asyncio=1.6.0
- networkx=3.4.2
- nomkl=3.0
- notebook=7.4.4
- notebook-shim=0.2.4
- numexpr=2.11.0
- numpy=1.26.4
- numpy-base=1.26.4
- openjpeg=2.5.2
- openldap=2.6.4
- openssl=3.0.17
- opentelemetry-api=1.30.0
- overrides=7.4.0
- packaging=25.0

- pandas=2.3.1
- pandoc=2.12
- pandocfilters=1.5.0
- parso=0.8.4
- pcre2=10.42
- pexpect=4.9.0
- pillow=11.3.0
- pip=25.1
- pixman=0.40.0
- platformdirs=4.3.7
- prometheus_client=0.21.1
- prompt-toolkit=3.0.43
- prompt_toolkit=3.0.43
- psutil=5.9.0
- pthread-stubs=0.3
- ptyprocess=0.7.0
- pure_eval=0.2.2
- pycparser=2.21
- pygments=2.19.1
- pyparsing=3.2.0
- pyqt=6.7.1
- pyqt6-sip=13.9.1
- pysocks=1.7.1
- python=3.10.18
- python-dateutil=2.9.0post0
- python-fastjsonschema=2.20.0
- python-json-logger=3.2.1
- python-tzdata=2025.2
- pytorch=2.6.0
- pytz=2025.2
- pyyaml=6.0.2
- pyzmq=26.2.0
- qtbase=6.7.3
- qtconsole=5.6.1
- qtdeclarative=6.7.3
- qtpy=2.4.1
- qtsvg=6.7.3
- qttools=6.7.3
- qtwebchannel=6.7.3
- qtwebsockets=6.7.3
- readline=8.3
- referencing=0.30.2
- requests=2.32.4
- rfc3339-validator=0.1.4
- rfc3986-validator=0.1.1
- rpds-py=0.22.3
- scikit-learn=1.7.1
- scipy=1.15.3

- seaborn=0.13.2
- send2trash=1.8.2
- setuptools=78.1.1
- sip=6.10.0
- six=1.17.0
- sleef=3.5.1
- sniffio=1.3.0
- soupsieve=2.5
- spirv-tools=2025.1
- sqlite=3.50.2
- stack_data=0.2.0
- sympy=1.13.3
- terminado=0.17.1
- threadpoolctl=3.5.0
- tinycss2=1.4.0
- tk=8.6.14
- tomli=2.2.1
- torchvision=0.15.2
- tornado=6.5.1
- traitlets=5.14.3
- typing-extensions=4.12.2
- typing_extensions=4.12.2
- tzdata=2025b
- unicodedata2=15.1.0
- urllib3=2.5.0
- wcwidth=0.2.13
- webencodings=0.5.1
- websocket-client=1.8.0
- wheel=0.45.1
- widgetsnbextension=4.0.13
- wrapt=1.17.0
- xcb-util=0.4.1
- xcb-util-cursor=0.1.5
- xcb-util-image=0.4.0
- xcb-util-keysyms=0.4.1
- xcb-util-renderutil=0.3.10
- xcb-util-wm=0.4.2
- xkeyboard-config=2.44
- xorg-libice=1.1.2
- xorg-libsm=1.2.6
- xorg-libx11=1.8.12
- xorg-libxau=1.0.12
- xorg-libxdmcp=1.1.5
- xorg-libxext=1.3.6
- xorg-libxfixes=6.0.1
- xorg-libxrandr=1.5.4
- xorg-libxrender=0.9.12
- xorg-libxshmfence=1.3.3

- xorg-libxxf86vm=1.1.6
- xorg-xorgproto=2024.1
- xz=5.6.4
- yaml=0.2.5
- zeromq=4.3.5
- zipp=3.21.0
- zlib=1.2.13
- zstd=1.5.6
